# Characterization of Target Gene Regulation by the Two Epstein-Barr Virus Oncogene LMP1 Domains Essential for B-cell Transformation

**DOI:** 10.1101/2023.04.10.536234

**Authors:** Bidisha Mitra, Nina Rose Beri, Rui Guo, Eric M. Burton, Laura A. Murray-Nerger, Benjamin E. Gewurz

## Abstract

The Epstein-Barr virus (EBV) oncogene latent membrane protein 1 (LMP1) mimics CD40 signaling and is expressed by multiple malignancies. Two LMP1 C-terminal cytoplasmic tail regions, termed transformation essential sites (TES) 1 and 2, are critical for EBV transformation of B lymphocytes into immortalized lymphoblastoid cell lines (LCL). However, TES1 versus TES2 B-cell target genes have remained incompletely characterized, and whether both are required for LCL survival has remained unknown. To define LCL LMP1 target genes, we profiled transcriptome-wide effects of acute LMP1 CRISPR knockout (KO) prior to cell death. To then characterize specific LCL TES1 and TES2 roles, we conditionally expressed wildtype, TES1 null, TES2 null or double TES1/TES2 null LMP1 alleles upon endogenous LMP1 KO. Unexpectedly, TES1 but not TES2 signaling was critical for LCL survival. The LCL dependency factor cFLIP, which plays obligatory roles in blockade of LCL apoptosis, was highly downmodulated by loss of TES1 signaling. To further characterize TES1 vs TES2 roles, we conditionally expressed wildtype, TES1 and/or TES2 null LMP1 alleles in two Burkitt models. Systematic RNAseq analyses revealed gene clusters that responded more strongly to TES1 versus TES2, that respond strongly to both or that are oppositely regulated. Robust TES1 effects on cFLIP induction were again noted. TES1 and 2 effects on expression of additional LCL dependency factors, including BATF and IRF4, and on EBV super-enhancers were identified. Collectively, these studies suggest a model by which LMP1 TES1 and TES2 jointly remodel the B-cell transcriptome and highlight TES1 as a key therapeutic target.

**Importance:** Epstein-Barr virus (EBV) causes multiple human cancers, including B-cell lymphomas. In cell culture, EBV converts healthy human B-cells into immortalized ones that grow continuously, which model post-transplant lymphomas. Constitutive signaling from two cytoplasmic tail domains of the EBV oncogene Latent Membrane Protein 1 (LMP1) is required for this transformation, yet there has not been systematic analysis of their host gene targets. We identified that only signaling from the membrane proximal domain is required for survival of these EBV-immortalized cells and that its loss triggers apoptosis. We identified key LMP1 target genes, whose abundance changed significantly with loss of LMP1 signals, or that were instead upregulated in response to switching on signaling by one or both LMP1 domains in an EBV-uninfected human B-cell model. These included major anti-apoptotic factors necessary for EBV-infected B-cell survival. Bioinformatics analyses identified clusters of B-cell genes that respond differently to signaling by either or both domains.

## Introduction

Epstein-Barr virus (EBV) is a gamma-herpesvirus that persistently infects most adults worldwide. EBV causes 200,000 cancers per year, including Burkitt lymphoma, Hodgkin lymphoma, post-transplant lymphoproliferative disease (PTLD), and HIV/AIDS associated lymphomas. EBV also causes a range of epithelial cell tumors, including gastric and nasopharyngeal carcinomas, as well as T and NK cell lymphomas(1). The key EBV oncogene latent membrane protein 1 (LMP1) is expressed in most of these tumors, where it drives growth and survival pathway signaling.

To colonize the B-cell compartment and establish lifelong infection, EBV uses a series of viral latency genome programs, in which different combinations of latency genes are expressed. These include six Epstein-Barr nuclear antigens (EBNA) and the membrane oncoproteins LMP1, LMP2A and LMP2B. LMP1 mimics aspects of signaling by the B-cell co-receptor CD40 (2-5), whereas LMP2A rewires surface B-cell immunoglobulin receptor signaling (6). All nine latency oncoproteins are expressed in the EBV B-cell transforming latency III program, which are expressed in immunoblastic lymphomas of immunosuppressed hosts. These include PTLD and primary central nervous system lymphoma. The latency II program is observed in EBV+ Hodgkin lymphoma, where the Reed-Sternberg tumor cells express EBNA1, LMP1 and LMP2A. Latency II is also frequently observed in T and NK cell lymphomas and in nasopharyngeal carcinoma. Host genome NF-κB activating mutations are frequently observed in EBV-negative Hodgkin lymphoma, but to a much lesser extent in EBV+ tumors, underscoring LMP1’s key role in activating growth and survival signaling (7).

LMP1 localizes to lipid rafts, where it signals constitutively in a ligand-independent fashion to activate NF-κB, MAP kinase, STAT3, PI3K, interferon and P62 pathways. LMP1 is comprised of a short N-terminal cytoplasmic tail, six transmembrane (TM) domains and a 200 residue C-terminal cytoplasmic tail (2-5). LMP1 TM domains drive homotypic aggregation, lipid raft association and constitutive signaling (8, 9). The LMP1 C-terminal tail functionally mimics signaling from activated CD40 receptors, to the point that the CD40 tail can essentially be replaced by that of LMP1 in transgenic mice studies. However, while CD40/LMP1 knockin mice had relatively normal B-cell development and evidence of intact CD40 function, including germinal center formation and class switch recombination, T-cell independent B-cell activation was also observed (10). These experiments suggest that the LMP1 C-terminal tail mimics CD40 signaling, but has also evolved additional functions.

Reverse genetic studies identified two LMP1 C-terminal cytoplasmic tail domains that are critical for EBV-mediated conversion of primary human B-cells into immortalized, continuously growing lymphoblastoid cell lines (LCL). Transformation effector site 1 (TES1), also called C-terminal activating region 1 (CTAR1), spans LMP1 residues 186-231. TES1/CTAR1 contains a PXQXT motif that engages tumor necrosis factor receptor associated factors (TRAFs). TES1 activates canonical NF-κB, non-canonical NF-κB, MAP kinase, PI3K and STAT3 pathways (3, 4, 11-16). TES2, which spans residues 351-386 and is also referred to as CTAR2, activates canonical NF-κB, MAPK, IRF7 and P62 pathways (3-5, 16-20). TRAF6 is critical for LMP1 TES2-driven canonical NF-κB, MAPK and p62 pathway activation (21-26). Canonical NF-κB signaling is critical for TES2/CTAR2 driven target gene regulation in a 293 cell conditional expression model (27). Signaling from a third LMP1 C-terminal tail region, CTAR3, activates JAK/STAT and SUMOylation pathways (28-30) potentially important *in vivo* but that are not essential for EBV-driven B-cell transformation (31). ChIP-seq analyses demonstrated a complex NF-κB binding landscape in LCLs, in which constitutive LMP1 signaling stimulates different combinations of the NF-κB transcription factors RelA, RelB, cRel, p50 and p52 to bind B-cell enhancers and promoters (32).

LMP1 is the only EBV oncogene that can independently transform rodent fibroblasts, driving anchorage independent growth and loss of contact inhibition (33-35). Notably, CTAR1 signaling is sufficient for LMP1-mediated fibroblast transformation, whereas CTAR2 was dispensable(36). LMP1 expression drives aberrant B-cell growth in transgenic B-cell models, particularly in combination with LMP2 upon disruption of cell mediated immunity (12, 37-40). While not critical for the first 8 days of EBV-driven B-cell outgrowth (41), LMP1 is critical for EBV-mediated conversion of primary human B-cells into immortalized LCLs (42, 43). A longstanding question has remained why TES1 and TES2 are each essential for EBV-mediated LCL establishment. Whether either or both are required for LCL survival is also unknown. Experiments using the EBV second site mutagenesis method (44) demonstrated that TES1 is critical for initiation of EBV-infected lymphoblastoid cell outgrowth (45). By contrast, TES2 is critical for long-term LCL growth, although TES2 null EBV infected lymphoblastoid cells could be propagated on epithelial feeders (45, 46). However, it remains incompletely understood the extent to which TES1 and TES2 play overlapping versus non-redundant roles.

While LMP1 B-cell target genes have been analyzed on small scales through qPCR and limited microarray analysis, unbiased genome-wide approaches have yet to be applied. Little is presently known about TES1 and TES2 shared versus non-redundant roles in transformed B-cells. To gain insights into LMP1 targets in the latency III LCL context, we therefore profiled transcriptome-wide changes in response to acute CRISPR LMP1 knockout (KO). These studies, performed at an early timepoint prior to apoptosis, identified that LMP1 strongly controls the LCL transcriptome, with expression levels of nearly 3400 host genes significantly altered by LMP1 KO. To then characterize specific LCL TES1 and TES2 roles, we conditionally expressed wildtype, TES1 null or TES2 null LMP1 rescue cDNAs at physiological levels upon endogenous LMP1 KO. This approach unexpectedly highlighted that signaling by TES1, but not TES2, is critical for LCL growth and survival. Loss of TES1 but not TES2 signaling rapidly triggered apoptosis, and strongly impaired expression of the LCL dependency factor cFLIP, which is required to block TNFα-driven apoptosis(25). Transcriptomic profiling further highlighted six clusters of LCL gene responses wildtype, TES1 or TES2 mutant LMP1, newly identifying independent, additive, or antagonistic roles in LCL target gene regulation. As multiple latency III genes often target the same host cell targets, we also constructed Burkitt B-cell models with conditional expression of wildtype, TES1 and/or TES2 null LMP1. These studies extended the LCL findings by further identifying shared versus distinct LMP1 roles in B-cell target gene regulation. Collectively, these studies highlight a complex landscape of TES1 and TES2 target gene regulation, in which each controls expression levels of large numbers of B-cell targets.

## Results

### CRISPR analysis of LCL LMP1 Target Genes

To characterize LMP1 target genes in the latency III context, we used CRISPR to knockout (KO) LMP1 in the well-characterized LCL GM12878, a Tier 1 Encode project cell line that we have used extensively for CRISPR analyses, and which we confirmed to have the latency III program (25, 47). GM12878 with stable Cas9 expression were transduced with lentivirus expressing a control single guide RNA (sgRNA) targeting a human genome intergenic region or LMP1. Immunoblot confirmed efficient LMP1 depletion by 48 hours post-puromycin selection of transduced LCLs (**Fig. 1A**). CRISPR LMP1 editing rapidly downmodulated the LMP1/NF-κB target genes TRAF1 and IRF4 and decreased non-canonical pathway processing of the p100 NF-κB precursor into the active p52 transcription factor subunit, suggesting successful on-target effects of LMP1 knockout (KO) (**Fig. 1A**). At this early timepoint post-CRISPR editing, LCLs remained viable (**Fig. S1A**). However, LMP1 KO triggers LCL growth arrest and cell death shortly thereafter. We therefore used this early 2-day post-puromycin selection timepoint to perform systematic RNAseq analyses of control vs LMP1 KO LCLs. At a multiple hypothesis testing adjusted p-value <0.05 and fold change of >2 cutoff, acute LMP1 KO significantly altered the levels of around 3400 host genes.

**Fig. 1.**
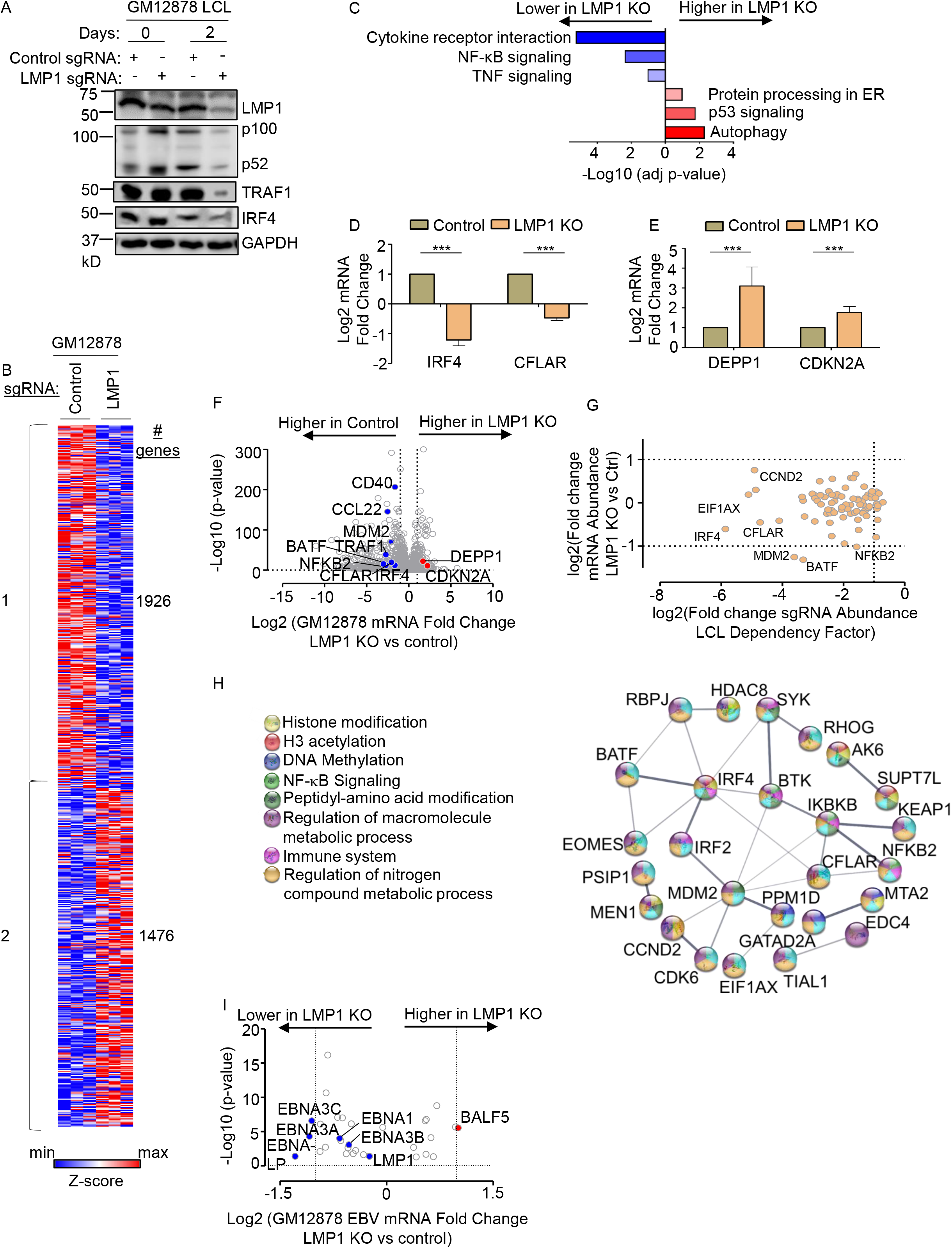
Characterization of LMP1 KO effects on GM12878 LCL target gene regulation. (A) Immunoblot analysis of whole cell lysates (WCL) from GM12878 LCLs transduced with lentiviruses that express control or LMP1 targeting single guide RNAs (sgRNAs). Transduced cells were puromycin selected for 0 vs 2 days, as indicated. Blots for LMP1, for LMP1 target genes TRAF1 and IRF4, and for LMP1-driven non-canonical NF-κB pathway p100/p52 processing are shown. Blots are representative of n=3 experiments. (B) K-means heatmap analysis of GM12878 LCLs transduced as in (A) with lentivirus expressing control or LMP1 sgRNA and puromycin selected for 48 hours. The heatmap depicts relative Z-scores in each row from n=3 independent RNAseq replicates, divided into two clusters. The Z-score scale is shown at bottom, where blue and red colors indicate lower versus higher relative expression, respectively. Two-way ANOVA P-value cutoff of <0.01 and >2-fold gene expression cutoffs were used. (C) Enrichr analysis of KEGG pathways most highly changed in GM12878 expressing control versus LMP1 sgRNA (LMP1 KO), as in (A). The X-axis depicts the -Log10 adjusted p-value (adj p-value) scale. The top three most enriched KEGG pathways are shown. (D) Abundances of two representative Cluster 1 genes from n=3 RNAseq analyses in cells with control vs LMP1 sgRNA. p-values were determined by one-sided Fisher’s exact test. *p<0.05, **p<0.01. (E) Abundances of two representative Cluster 2 genes from n=3 RNAseq analyses in cells with control vs LMP1 sgRNA. p-values were determined by one-sided Fisher’s exact test. ***p<0.001. (F) Volcano plot analysis of host transcriptome-wide GM12878 genes differentially expressed in cells with control versus LMP1 sgRNA expression, as in (B), using data from n=3 RNAseq datasets. (G) Scatter plot cross comparison of log2 transformed fold change mRNA abundances in GM12878 expressing LMP1 vs. control sgRNA (Y-axis) versus log2 transformed fold change abundances of sgRNAs at Day 21 versus Day 1 post-transduction of GM12878 LCLs in a genome-wide CRISPR screen (25) (X-axis). (H) String analysis of genes shown in (G). Pathway identifiers for each gene and interaction are colored coded. (I) Volcano plot analysis of EBV mRNA values in GM12878 expressing LMP1 vs control sgRNAs, as in (B). p-value <0.05 and >2-fold change mRNA abundance cutoffs were used.

Genes differentially expressed in LMP1 KO vs control cells could be broadly characterized into two k-means clusters, in which LMP1 KO either downregulated 1,476 or upregulated 1926 host genes (**Fig. 1B, Table S7**). Kyoto Encyclopedia of Genetic Elements (KEGG) pathway Enrichr analysis (48) identified that cytokine receptor signaling, NF-κB signaling and tumor necrosis factor (TNF) signaling as enriched amongst genes rapidly downmodulated by LMP1 KO (**Fig. 1C**). As examples of cluster 1 NF-κB target genes, interferon regulatory factor 4 (IRF4) and CFLAR, which encodes c-FLIP, were strongly downmodulated by LMP1 KO (**Fig. 1D**). By contrast, KEGG highlighted that autophagy, p53 signaling and protein-processing in the endoplasmic reticulum as enriched amongst cluster 2 genes (**Fig. 1C**). As examples from these enriched pathways, LMP1 KO highly induced expression of the autophagy suppressor DEPP1 and the p53 target and tumor suppressor cyclin dependent kinase inhibitor 2A (CDKN2A) (**Fig. 1D-F**). Quantitative real-time PCR (qRT-PCR) validated RNAseq results for the CCL22, EBI3 and IRF4 mRNAs, each of which were significantly downmodulated by LMP1 KO (**Fig. S1B-D**).

We next integrated our RNAseq dataset with published CRISPR analysis of host dependency factors essential for EBV+ LCL, but not Burkitt B-cell proliferation (25), to gain insights into key LMP1 roles in LCL growth and survival. This analysis identified that mRNA abundances of 37 of the 87 CRISPR-defined LCL selective dependency factors significantly changed upon LMP1 KO, suggesting multiple LMP1 roles in support of LCL survival (**Fig. 1G and S1E**). Of these, it is notable that multiple key suppressors of LCL intrinsic and extrinsic apoptotic pathways were rapidly lost upon LCL LMP1 KO. For instance, our published CRISPR analyses highlighted non-redundant roles for the transcription factors IRF4 and BATF in blockade of the intrinsic apoptosis pathway and for CFLAR-encoded cFLIP in extrinsic apoptosis pathway inhibition (25), each of whose mRNAs rapidly decreased upon LCL LMP1 KO. Likewise, LMP1 KO strongly downmodulated expression of MDM2, an LCL-selective dependency factor (25) that targets p53 for proteasomal degradation and that prevents LCL p53-dependent apoptosis(49). Furthermore, NF-κB blockade triggers LCL apoptosis(50) and the LCL dependency factor NFKB2, which encodes the NF-κB transcription factor subunit p52, was also highly downmodulated by LMP1 KO (**Fig. 1F-G**). STRING network analysis also underscored that each of these assemble into a network with 23 other LMP1-regulated LCL dependency factors (**Fig. 1H**).

Since LMP1 is highly expressed in Hodgkin lymphoma Reed-Sternberg tumor cells, we next analyzed effects of LMP1 KO on Hodgkin lymphoma KEGG pathway genes (**Fig. S1F**). Interestingly, LMP1 KO strongly downmodulated expression of the T-cell tropic chemokines CCL22 and CCL17, consistent with several prior reports linking LMP1 to their expression (51, 52). These findings raise the possibility that LMP1-driven chemokine expression may contribute to the striking enrichment of T-cells characteristic of the Hodgkin Reed-Sternberg microenvironment. However, volcano plot analysis also highlighted that LMP1 KO increased expression of CD274, which encodes the checkpoint inhibitor PD-L1, further implicating LMP1 in T-cell regulation.

Given widespread effects of LMP1 KO on LCL host gene expression, we next characterized effects of LMP1 KO on viral latency III genes. Mapping of RNAseq reads onto the GM12878 EBV transcriptome identified that LMP1 depletion significantly downmodulated mRNAs encoding EBNA3A, 3C and EBNA-LP, though interestingly not those encoding EBNA2 or EBNA3B (**Fig. 1I**). While it has been reported that LMP1 regulates its own mRNA expression (53, 54), we did not observe changes in LMP1 mRNA abundance upon LMP1 CRISPR KO. We note that CRISPR editing often results in insertions or deletions, causing functional protein knockout without necessarily changing mRNA levels of the edited gene. However, it is plausible a compensatory response to LMP1 knockout occurred on the mRNA level at this early timepoint, potentially balancing loss of NF-κB induced LMP1. Taken together, our RNAseq analyses raise the possibility that secondary effects of LMP1 KO on Epstein-Barr nuclear antigens may also contribute to changes in the host transcriptome and cell death upon LMP1 KO.

### TES1 but not TES2 signaling is critical for LCL survival

While TES1 and TES2 signaling are each critical for B-cell transformation, it has remained unknown whether either or both are necessary for proliferation of fully transformed LCLs. Likewise, knowledge has remained incomplete about shared versus non-redundant TES1 and TES2 roles in LCL host gene regulation. To gain insights into these key questions, we engineered Cas9+ GM12878 LCLs with conditional expression of wildtype (WT) LMP1, or with well characterized point mutants that abrogate signaling from the TES1 TRAF binding domain (TES1m, residues 204PQQAT208 -> **A**Q**A**AT), from TES2 (TES2m, 384YYD386 -> ID) (46, 55, 56) (**Fig. 2A**). A silent mutation in the CRISPR protospacer adjacent motif (PAM) was used to abrogate CRISPR editing of these LMP1 rescue cDNA constructs. For cross-comparison, we also established conditional TES1/TES2 double mutant (DM) cell lines with both mutations, to profile responses to other LMP1 regions, potentially including CTAR3 or unfolded protein responses induced by LMP1 induction (57, 58) (**Fig. 2A**). LCLs were then transduced with lentivirus expressing a control sgRNA targeting a human intergenic region or LMP1. Conditional LMP1 expression was then induced by addition of 400 ng/ml doxycycline, such that the rescue cDNA was induced as endogenous EBV-encoded LMP1 was depleted. We confirmed similar levels of LMP1 expression across this series and achieved similar LMP1 levels as in unedited GM12878 LCLs (**Fig. 2B**). Importantly, we validated that WT LMP1 rescued physiological levels of LMP1 target TRAF1 expression and p100/p52 processing (**Fig. 2B**). TES1 is responsible for the majority of LMP1-mediated TRAF1 induction and p100/p52 processing(55, 59) and as expected, conditional TES1m and DM expression failed to rescue physiological levels of TRAF1 or p100/p52 processing in GM12878 with endogenous LMP1 KO. By contrast, TES2m induced levels of TRAF1 and p100/p52 processing in LMP1 KO levels approaching those in unedited GM12878 (**Fig. 2B**), validating our LCL LMP1 KO/rescue system.

**Fig. 2.**
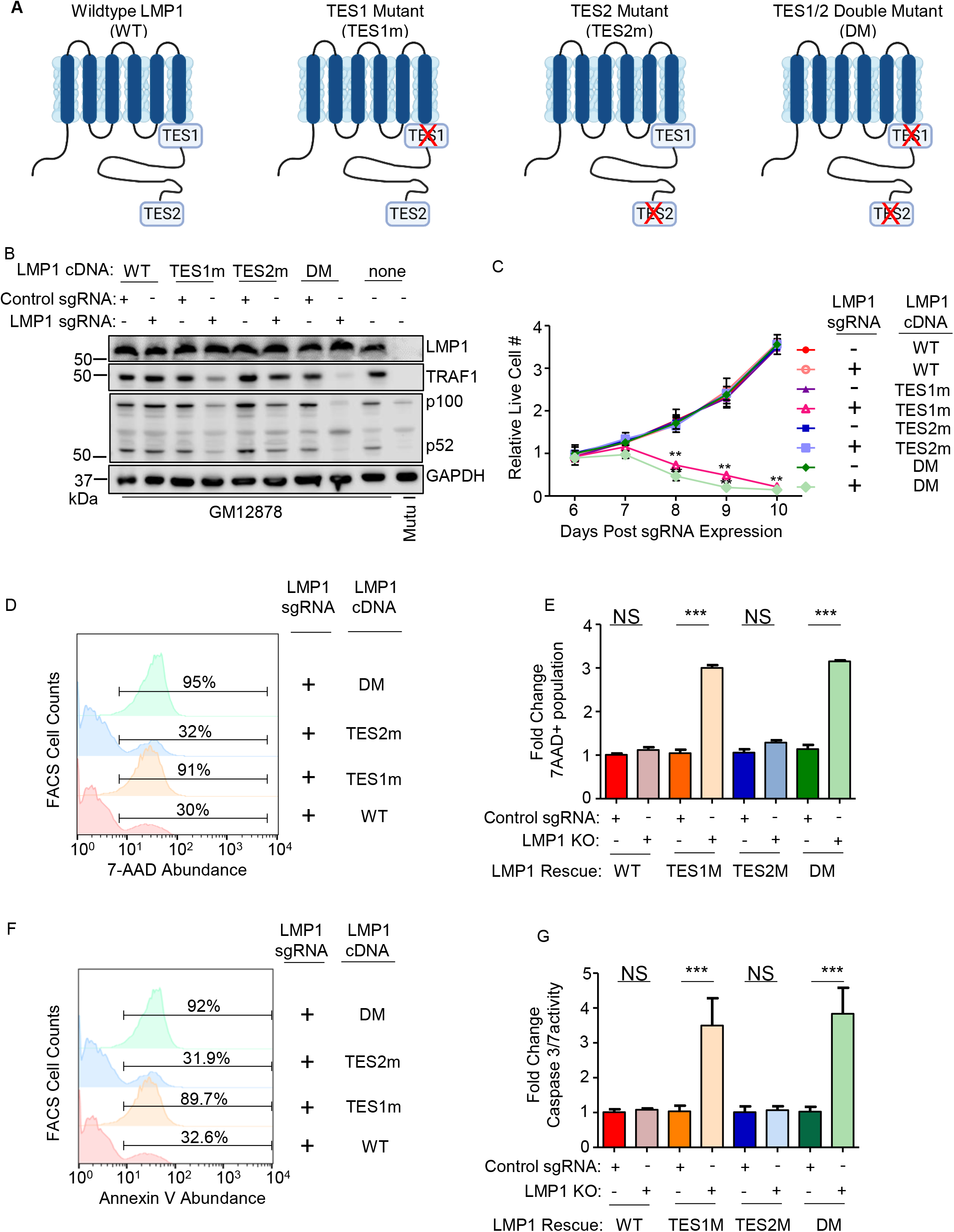
Loss of TES1 but not TES2 signaling triggers LCL apoptosis. (A) Schematic diagram of LMP1 WT with TES1 and TES2 domains highlighted. Wildtype (WT) or point mutants abrogated for signaling from TES1 (TES1m), TES2 (TES2m) or double TES1/TES2 mutant (DM) are shown. (B) Immunoblot analysis of WCL from GM12878 LCLs that expressed control or LMP1 sgRNAs and puromycin selected for 3 days, then induced for expression with the indicated LMP1 rescue cDNA construct for 6 days. Blots are representative of n = 3 experiments. (C) Growth curve analysis of GM12878 LCLs at the indicated day post expression of control or LMP1 sgRNAs and the indicated LMP1 WT, TES1m, TES2m or DM rescue cDNA. Shown are mean ± SD from n=3 independent experiments. **p<0.01. (D) FACS analysis of 7-AAD vital dye uptake in GM12878 on day 7 post-expression of LMP1 sgRNAs and the indicated LMP1 rescue cDNA. Shown are percentages of 7-AAD+ cells within the indicated gates. Representative of n=3 experiments. (E) Mean ± SD of fold change 7-AAD values from n=3 independent experiments of GM12878 with the indicated control or LMP1 sgRNA and rescue cDNA expression, as in (D). Values in GM12878 with control sgRNA and no LMP1 rescue cDNA were set to 1. (F) FACS analysis of plasma membrane annexin V abundance in GM12878 on day 7 post-expression of control or LMP1 sgRNAs and the indicated LMP1 rescue cDNA. Shown are percentages of 7-AAD+ cells within the indicated gates. Representative of n=3 experiments. (G) Mean ± SD of fold-change caspase 3/7 activity levels, as determined by caspase 3/7 Glo assay, from n=3 independent experiments of GM12878 with the indicated control or LMP1 sgRNA and rescue cDNA expression. Values in GM12878 with control sgRNA and no LMP1 rescue cDNA were set to 1.

We next tested the effects of LMP1 WT, TES1m, TES2m or DM rescue cDNA expression on GM12878 proliferation. Whereas signaling from both TES1 and TES2 are required for EBV-driven primary human B-cell growth transformation, we unexpectedly found that LCLs require signaling only from TES1 for growth and survival: LMP1 KO LCLs with WT vs TES2m rescue cDNA proliferated indistinguishably. By contrast, LCLs with TES1m or DM rescue cDNA expression failed to proliferate (**Fig. 2C**). To characterize this unexpected result further, we next measured effects of endogenous LMP1 KO and rescue LMP1 cDNA expression on LCL survival. Consistent with our growth curve analysis, LMP1 KO LCLs with TES1m or DM rescue cDNA exhibited widespread cell death, as judged by uptake of the vital dye 7aminoactinomycin D (7-AAD) by FACS analysis (**Fig. 2D-E**). Consistent with apoptosis as the cell death pathway triggered by loss of TES1 signaling in LCLs, levels of Annexin V and executioner caspase 3/7 activity were significantly higher in LMP1 KO GM12878 with TES1m or DM than with WT or TES2m rescue cDNA expression (**Fig. 2F-G and S2**). Taken together, these data newly suggest that TES1 signaling is necessary for LCL growth and survival in a manner that is not redundant with TES2, and that cannot be rescued by TES2 signaling alone.

### Identification of TES1 versus TES2 roles in LCL gene regulation

To gain insights into overlapping versus non-redundant TES1 and TES2 LCL roles, we performed biological triplicate RNAseq analyses to cross-compare GM12878 transcriptomes at day 6 post endogenous LMP1 KO and with doxycycline-induced WT, TES1m or TES2m rescue cDNA expression. We selected this early timepoint as it is just prior to the divergence of the growth curves (**Fig. 2C, Table S8-9**) and the onset of apoptosis. K-means analysis identified six clusters in which host gene expression differed between LMP1 KO LCLs with WT, TES1m or TES2m cDNA rescue. KEGG analysis highlighted pathways most highly enriched in each cluster (**Fig. 3A**). Notably, apoptosis pathway genes were the most highly enriched in cluster 4 genes, which were expressed at lower levels in cells with TES1m than with WT or TES2m rescue cDNA expression, suggesting that TES1 signaling may induce their expression. Apoptosis genes were also enriched amongst cluster 5 genes, where levels were lower in cells with TES2m expression, suggesting that TES2 signaling may induce their expression (**Fig. 3A**).

**Fig. 3.**
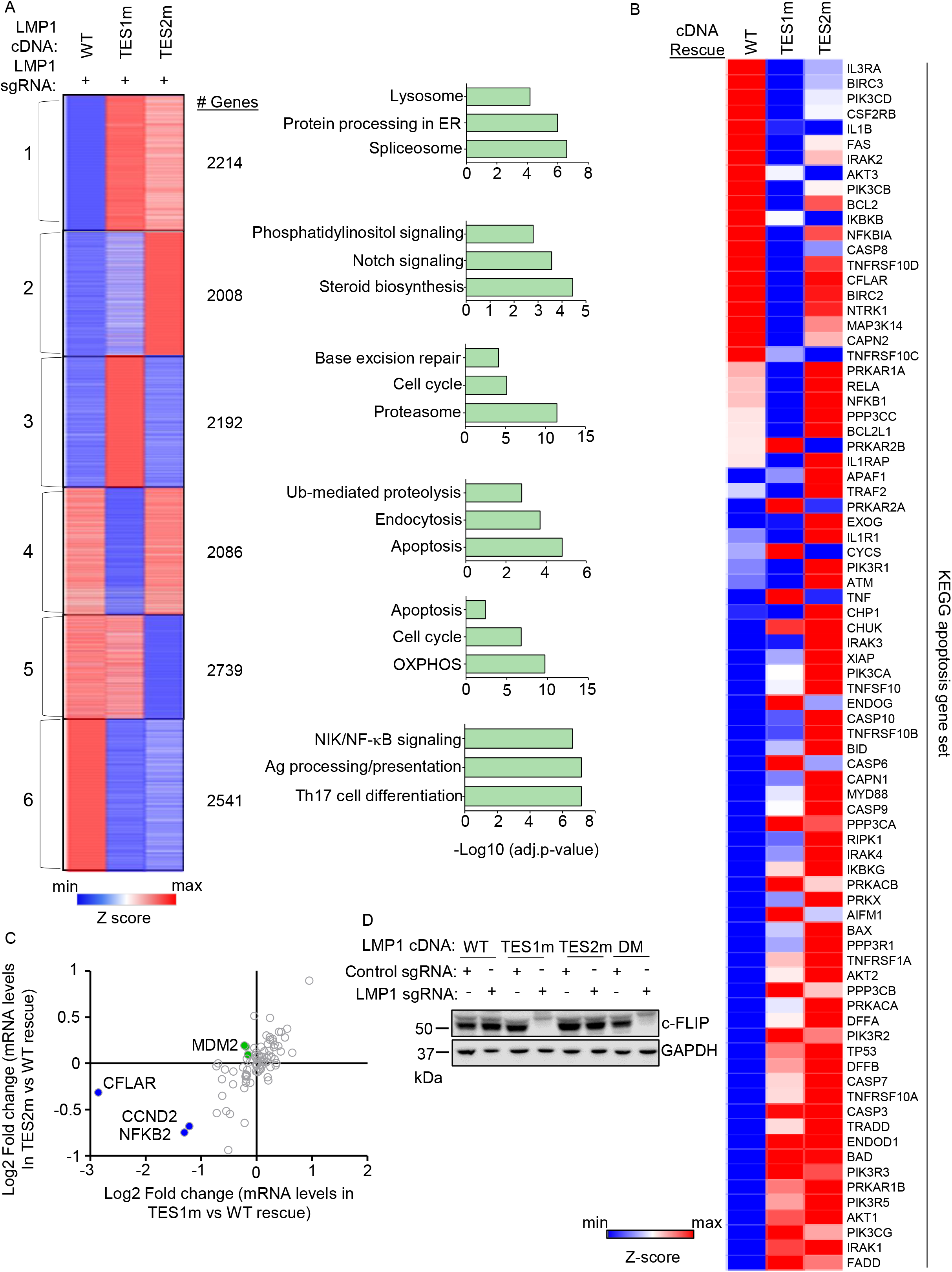
Characterization of host genome-wide TES1 vs TES2 LCL target genes. (A) RNAseq K-means heatmap analysis of GM12878 LCLs transduced with lentivirus expressing LMP1 sgRNA and induced for WT, TES1m or TES2m rescue cDNA expression for 6 days. The heatmap depicts relative Z-scores in each row from n=3 independent RNAseq datasets, divided into six clusters. The Z-score scale is shown at bottom, where blue and red colors indicate lower versus higher relative expression, respectively. Two-way ANOVA P-value cutoff of <0.05 and >2-fold gene expression cutoffs were used. The top three most highly enriched KEGG pathways amongst genes within each cluster are shown at right. (B) Heatmap analysis of KEGG apoptosis pathway gene relative row Z-scores from RNAseq analysis as in (A). The Z-score scale is shown at bottom, where blue and red colors indicate lower versus higher relative expression, respectively. Two-way ANOVA P-value cutoff of <0.05 and >2-fold gene expression cutoffs were used. (C) Scatter plot analysis cross comparing log2 transformed fold change of LCL dependency factor mRNA abundances in GM12878 expressing LMP1 sgRNA together with TES2 mutant versus wildtype cDNA rescue (Y-axis) and TES1 mutant versus wildtype cDNA rescue (X-axis) from triplicate RNAseq datasets, as in (A). This analysis highlighted that CFLAR and to a lesser extent NFKB2 and CCND2 mRNAs were more highly downmodulated by TES1m than TES2m rescue, relative to levels in cells with WT LMP1 rescue. Shown are genes differentially regulated by >2 fold with either TES1m or TES2m rescue, relative to levels with WT LMP1 rescue. (D) Immunoblot analysis of c-FLIP and load control GAPDH expression in WCL from GM12878 LCLs with the indicated control or LMP1 sgRNA and LMP1 rescue cDNA expression. Representative of n=3 experiments.

Given this apoptosis signal, we next analyzed the KEGG apoptosis gene set responses to WT, TES1m or TES2m cDNA rescue (**Fig. 3B**). Interestingly, a cluster of genes were more highly expressed in cells with WT and TES2m than with TES1m expression, including the anti-apoptotic genes CFLAR, BCL2 and BIRC3, which encodes the cIAP2 ubiquitin ligase, which counteracts TNF-driven cell death. BCL2 and BIRC3 were not defined as LCL dependency factors by genome-wide CRISPR analysis, whereas CLFAR was (25). Intriguingly, while a small number of the 87 LCL dependency factors (25) were more highly downmodulated in LMP1 KO LCLs with TES1 rescue than with TES2 rescue, CFLAR was the LCL dependency factor most highly depleted from with TES1m rescue (nearly 8-fold) (**Fig. 3C, S3A**). By contrast, CFLAR was only mildly depleted (<0.5 fold) in cells with TES2m rescue (**Fig. 3C**). We validated that CFLAR-encoded c-FLIP was highly downmodulated on the protein level in LMP1 KO LCLs with TES1m or DM LMP1 rescue, but not in LCLs with WT or TES2m rescue at an early timepoint prior to cell death (**Fig. 3D**). The LCL dependency factors NFKB2 and CCND2 were also more highly downmodulated in LMP1 KO cells with TES1 than TES2 cDNA rescue but not to the same extent as CFLAR (**Fig. 3C**), suggesting that their loss may not be responsible for apoptosis in the absence of TES1 signaling. Collectively, these analysis underscore distinct TES1 versus TES2 signaling roles in control of LCL apoptosis pathway gene expression.

To gain further insights into potential TES1 vs TES2 roles in regulation of genes with relevance to Hodgkin Reed-Sternberg cells, we analyzed KEGG Hodgkin lymphoma pathway gene expression in LMP1 KO GM12878 rescued with WT, TES1m or TES2m LMP1. This analysis highlighted that many KEGG Hodgkin lymphoma pathway genes are jointly induced by TES1 and TES2 signaling in LCLs, including CCL22, BCL3, cRel, IRF4, STAT3, STAT6 and CD70, each of which have prominent roles in Hodgkin lymphoma pathogenesis (**Fig. S3B**). We also confirmed the transcript levels of CCL22, EBI3 and IRF4 through qRT-PCR and the results were concordant with our RNA-seq analysis where TES1 and TES2 are jointly responsible in the induction of the expression of these genes (**Fig. S3 C-D**). Interestingly, the Hodgkin lymphoma therapeutic target CD27 was expressed at lower level with TES1m rescue but more highly expressed in cells with TES2m rescue, suggesting that TES1 and TES2 signaling may jointly balance its expression.

On the transcriptome-wide level, multiple well characterized LMP1 target genes were more highly expressed in LCLs rescued with WT LMP1 than with TES1m cDNA, establishing these as key TES1 LCL target genes. These included CD40, TRAF1, EBI3 and ICAM1 (**Fig. 4A**), which was previously established as a TES1 target genes in studies of cell lines with LMP1 over-expression, including BL-41 Burkitt and BJAB diffuse large B-cell lymphoma models(55). Notably, this approach also newly suggests a large number of B-cell targets whose upregulation or downregulation is dependent on TES1 signaling. These include CFLAR and TLR6 (which encodes Toll-like receptor 6), which were significantly more highly expressed in LMP1 KO LCLs with WT LMP1 cDNA rescue. By contrast, the mRNA encoding the histone loader DAXX, which can serve as an epigenetic suppressor of EBV gene expression (60, 61) and DNA damage pathway TP53 (which encodes p53), were expressed at considerably higher in LMP1 KO LCLs with TES1m than WT LMP1 rescue cDNA (**Fig. 4A**). This result suggests that TES1 may repress their expression. Enrichr analysis of genes more highly expressed with WT LMP1 rescue highlighted TNF and NF-κB signaling as enriched KEGG pathways, whereas p53 signaling and apoptosis were amongst the pathways most highly enriched in genes more highly expressed with TES1m rescue (**Fig. 4B**).

**Fig. 4.**
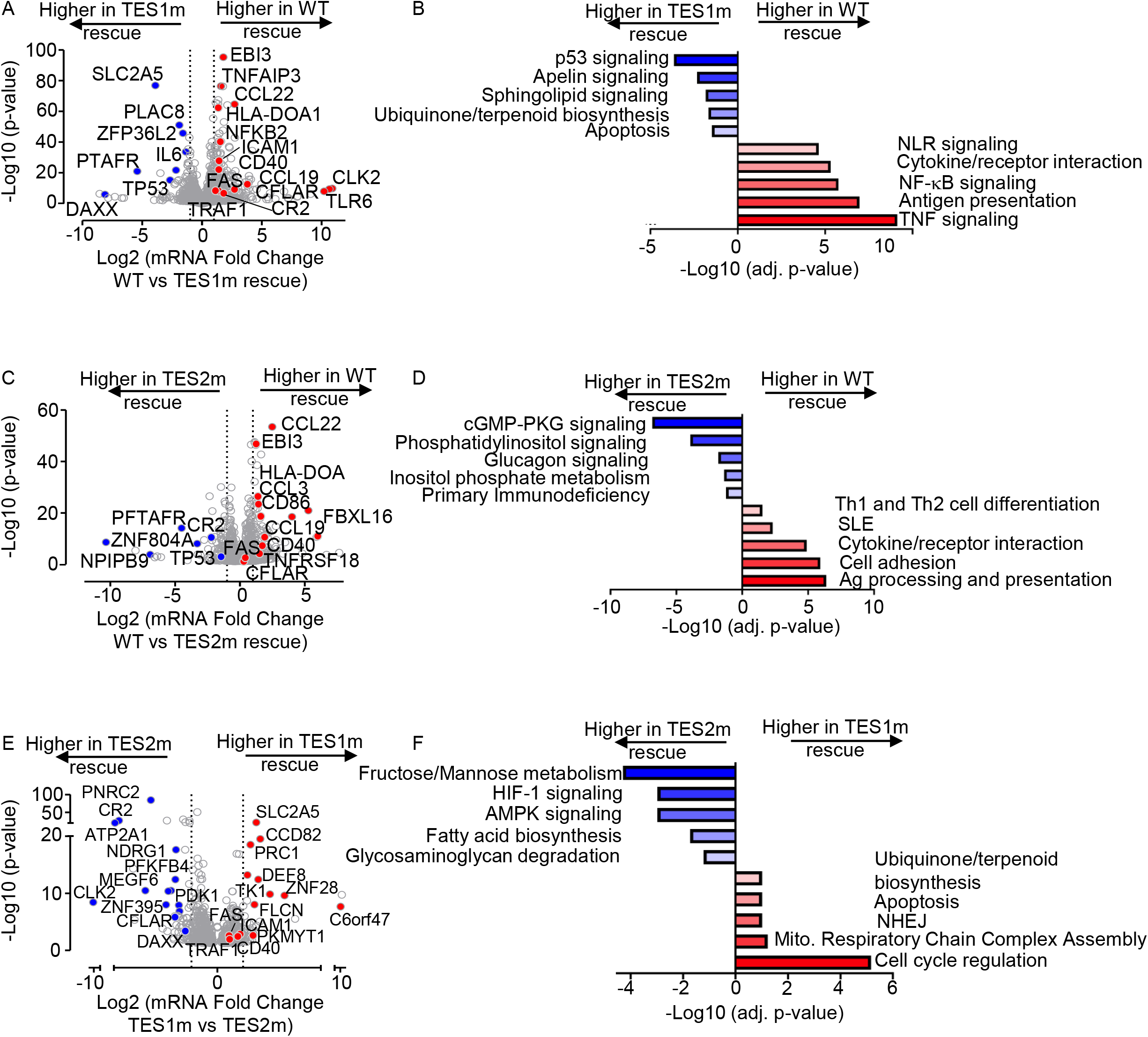
Characterization of LCL pathways targeted by TES1 vs TES2 signaling. (A) Volcano plot analysis of host transcriptome-wide GM12878 genes differentially expressed in LMP1 KO GM12878 with WT vs TES1 mutant cDNA rescue. Higher X-axis fold changes indicate higher expression with WT LMP1 rescue, whereas lower X-axis fold changes indicate higher expression with TES1m rescue. Data are from n=3 RNAseq datasets, as in **Fig. 3**. (B) Enrichr analysis of KEGG pathways most highly enriched in RNAseq data as in (A) amongst genes more highly expressed in LMP1 KO GM12878 with WT than TES1m rescue (red) vs amongst genes more highly expressed with TES1m than WT rescue (blue). (C) Volcano plot analysis of host transcriptome-wide GM12878 genes differentially expressed in LMP1 KO GM12878 with WT vs TES2 mutant cDNA rescue. Higher X-axis fold changes indicate higher expression with WT LMP1 rescue, whereas lower X-axis fold changes indicate higher expression with TES2m rescue. Data are from n=3 RNAseq datasets, as in **Fig. 3**. (D) Enrichr analysis of KEGG pathways most highly enriched in RNAseq data as in (C) amongst genes more highly expressed in LMP1 KO GM12878 with WT than TES2m rescue (red) vs amongst genes more highly expressed with TES2m than WT rescue (blue). (E) Volcano plot analysis of host transcriptome-wide GM12878 genes differentially expressed in LMP1 KO GM12878 with TES1 vs TES2 mutant cDNA rescue. Higher X-axis fold changes indicate higher expression with TES1m rescue, whereas lower X-axis fold changes indicate higher expression with TES2m rescue. Data are from n=3 RNAseq datasets, as in **Fig. 3**. (F) Enrichr analysis of KEGG pathways most highly enriched in RNAseq data as in (E) amongst genes more highly expressed in LMP1 KO GM12878 with TES1m than TES2m rescue (red) vs amongst genes more highly expressed with TES2m than TES1m rescue (blue).

Volcano plot and KEGG pathway analysis highlighted LCL genes differentially expressed in LMP1 KO LCLs with WT vs TES2m rescue (**Fig. 4C-D**). KEGG pathways enriched amongst genes more highly expressed with WT LMP1 rescue again included antigen presentation and cytokine/receptor interaction, but also included systemic lupus erythematosus (SLE). Pathways enriched amongst genes more highly expressed with TES2m rescue instead included cyclic GMP protein kinase G (cGMP-PKG) signaling and phosphatidylinositol signaling (**Fig. 4D**). Notably, TP53 (which encodes p53) was also more highly expressed in LMP1 KO LCLs with TES2m than WT LMP1 rescue, suggesting that both TES1 and 2 signaling regulate its expression. By comparison, CFLAR expression was similar in LMP1 KO LCLs with WT and TES2m rescue, further establishing it as a TES1 target in LCLs (**Fig. 4C**).

Direct cross-comparison of genes in LCLs with TES1m vs TES2m rescue further identified roles of TES1 versus TES2 signaling on LCL target gene expression. The oncogenic kinase CLK2, which has roles in splicing regulation, was the host gene most highly expressed in LMP1 KO with TES2m rescue than in cells with TES1m rescue, newly indicating that it is strongly induced by TES1 or strongly inhibited by TES1 (**Fig. 4E**). Enrichr analysis indicated that multiple KEGG metabolism pathways were the most highly enriched in cells with TES2m rescue, including fructose/mannose metabolism, HIF1 signaling and AMPK signaling (**Fig. 4E-F**). In support, the glycolytic enzyme PFKFB4 and the kinase PDK1, which regulates flux of glycolytic products to mitochondrial metabolism pathways at the level of pyruvate, were more highly expressed with TES2m rescue, suggesting that they are either driven by TES1 or repressed by TES2 signaling (**Fig. 4F**). Cell cycle regulation was the KEGG pathway most enriched amongst genes more highly expressed with TES1m rescue. The cyclin dependent kinase (CDK) substrate and cytokinesis regulator PRC1, as well as the CDK1 kinase and mitosis regulator PKMYT1, were amongst the genes most highly differentially expressed in TES1m rescue (Fig. **4E-F**), suggesting that TES2 drives or that TES1 instead inhibits their expression.

### B-cell genes induced by conditional expression LMP1 in EBV-negative Burkitt cells

As a complementary approach to our loss-of-function CRISPR KO LCL analyses, we next profiled B-cell responses to conditional LMP1 expression. A goal of this approach was to identify LMP1-specific effects on host gene expression, since LMP1 KO significantly altered expression of several EBV latency III genes. Furthermore, EBNA and LMP latency III oncoproteins often jointly target host genes. Therefore, to study LMP1-specific effects in isolation of other latency III genes, we engineered EBV-negative Akata and BL-41 Burkitt B-cell lines with doxycycline-inducible WT, TES1m, TES2m or DM LMP1 alleles. The Akata cell line was originally established from a human EBV+ Burkitt tumor (62), but an EBV-negative subclone that spontaneously lost the viral genome was isolated shortly thereafter (63), which we used for these studies. Similarly, the BL-41 cell line was established from an EBV-negative human Burkitt lymphoma tumor (64). BL-41 were used for early microarray analysis of latency III or LMP1 effects on a subset of human genes (50). We validated that WT and point mutant LMP1 were expressed to similar extents across the panel. As expected, TES1m and DM exhibited impaired non-canonical NF-κB pathway activation, as judged by p100:p52 processing (**Fig S4A-B**). LMP1 signaling was also validated by FACS analysis of ICAM-1 and Fas upregulation. Consistent with a published study (55), TES1 signaling more strongly induced ICAM-1 and Fas in both Burkitt cell lines, even though BL-41 had somewhat higher basal NF-κB activity than Akata, as judged by Fas and ICAM-1 levels in uninduced cells (**Fig. S4C-J**).

We then profiled effects of conditional WT, TES1m, TES2m or DM expression for 24 hours on the Akata transcriptome using biological triplicate RNAseq datasets. K-means heatmap analysis with n=6 clusters revealed strikingly distinct patterns of host gene responses to WT, TES1m, TES2m and DM LMP1 signaling (**Fig. 5A, Table S1-3**). Cluster 1 genes were highly upregulated by WT LMP1, to a lesser extent by TES2m (in which only TES1 signals), and more modestly by TES1m (in which only TES2 signals). This result suggests that TES1 signaling contributes more strongly than TES2 to their expression. Notably, CFLAR was a cluster I gene target, consistent with our finding that TES1 drives CFLAR expression in LCLs, as was the interferon stimulated gene IFIT1 (**Fig. 5B**). KEGG pathways enriched amongst Cluster 1 genes included TLR signaling, chemokine signaling, IFN signaling and NLR signaling (**Fig. 5C**). Cluster 1 also contained well described LMP1 target genes, including TRAF1, which we validated by immunoblot (**Fig. S4A**), consistent with a prior study(55). Notably, MAP3K7, which encodes the kinase TAK1 is also a Cluster 1 gene. Since TAK1 is critical for TES2/canonical NF-κB and MAP kinase signaling (26), this result suggests an important mechanism of cross-talk between TES1 and TES2. Likewise, the Cluster I gene IRF7 binds to and is activated by TES2 (65-68), again suggesting cross-talk between LMP1 pathways. STAT1 and STAT3 are also Cluster 1 genes, raising the question of whether these STATs may drive interferon stimulated gene induction downstream of LMP1.

**Fig. 5.**
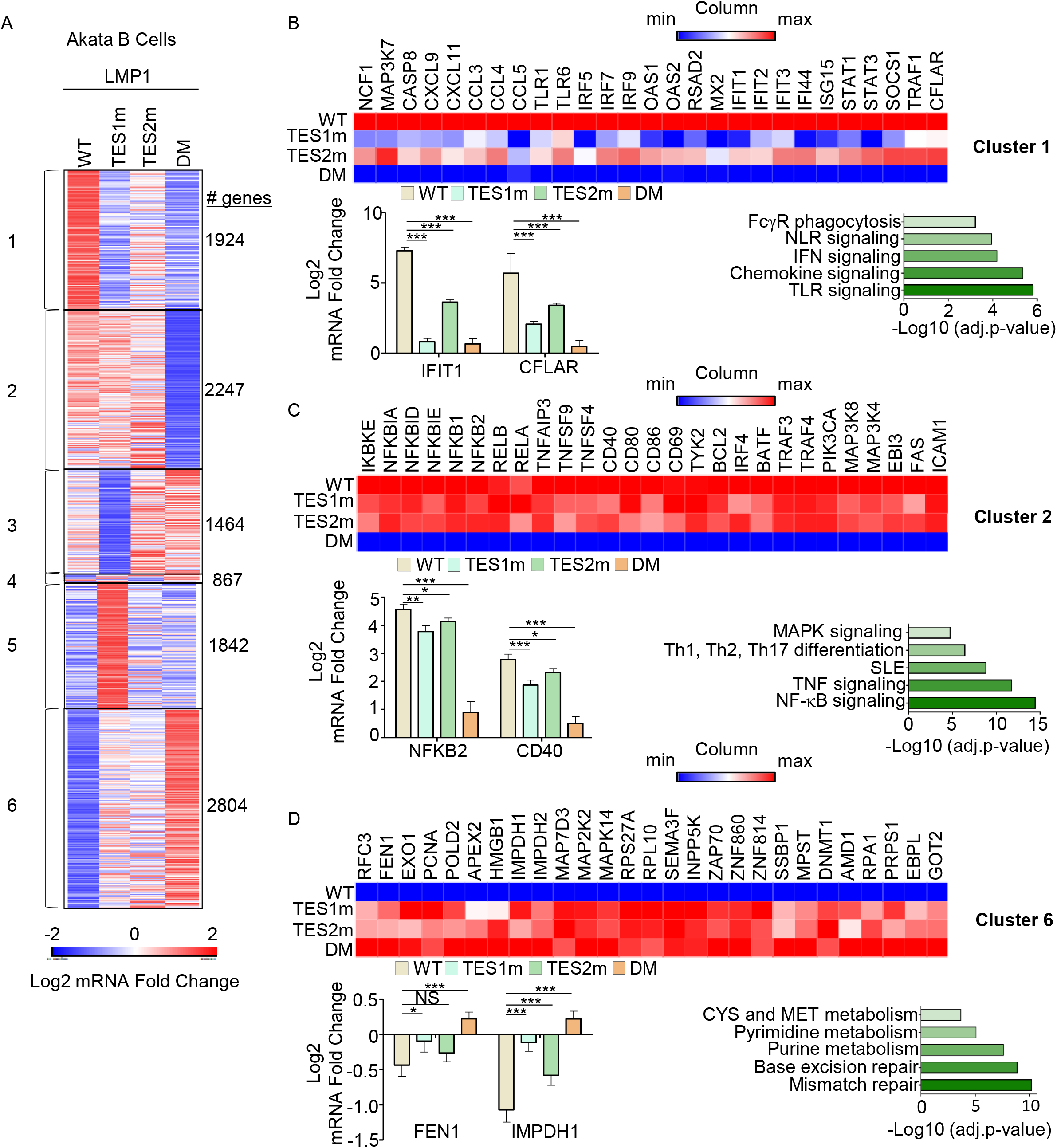
Characterization of host genome-wide Akata B-cell LMP1 target genes. (A) K-means heatmap analysis of RNAseq datasets from n=3 replicates generated in EBV-Akata Burkitt cells with conditional LMP1 WT, TES1m, TES2m or DM expression induced by 250 ng/ml doxycycline for 24 hours. The heatmap visualizes host gene Log2 Fold change across the four conditions, divided into six clusters. A two-way ANOVA P value cutoff of <0.01 and >2-fold gene expression were used. # of genes in each cluster is indicated at right. (B) Heatmaps of representative Cluster 1 differentially regulated genes (top), with column maximum (max) colored red and minimum (min) colored blue, as shown by the scalebar. Also shown are expression values of two representative Cluster 1 genes (lower left) and Enrichr analysis of KEGG pathways significantly enriched in Cluster 1 gene sets (lower right). p-values were determined by one-sided Fisher’s exact test. ***p<0.001. (C) Heatmaps of representative Cluster 2 differentially regulated genes (top), as in (B). Also shown are expression values of two representative Cluster 2 genes (lower left) and Enrichr analysis of KEGG pathways significantly enriched in Cluster 2 gene sets (lower right). p-values were determined by one-sided Fisher’s exact test. *p<0.05, **p<0.01, ***p<0.001. (D) Heatmaps of representative Cluster 6 differentially regulated genes (top), as in (B). Also shown are expression values of two representative Cluster 6 genes (lower left) and Enrichr analysis of KEGG pathways significantly enriched in Cluster 6 gene sets (lower right). p-values were determined by one-sided Fisher’s exact test. *p<0.05, ***p<0.001.

Cluster 2 genes were induced by WT, TES1m or TES2m LMP1 to a similar extent (**Fig. 5C**), suggesting that they redundantly respond to TES1 or TES2 signaling. GO analysis highlighted enrichment of NF-κB signaling in this cluster, and which included mRNAs encoding four NF-κB transcription factor subunits, as well as the NF-κB induced inhibitors IκBα, IκBζ and IκBε. mRNA fold changes for *NFBK2*, which encodes non-canonical pathway NF-κB p52 transcription factor are shown in **Fig. 5C** and are consistent with our LCL rescue analysis, which identified similarly important roles for both TES1 and TES2 in support of NFKB2 expression (**Fig. 3C**). Consistent with a prior study (55), Fas and ICAM-1 are Cluster 2 genes similarly induced on the mRNA level by TES1m and TES2m, though interestingly, plasma membrane ICAM-1 levels were lower in cells expressing TES1m (**Fig. S4C-F**). This result raises the possibility that TES1 signaling may play a role in ICAM-1 post-transcriptional regulation and/or trafficking.

Cluster 3 genes were expressed at lower levels in cells expressing TES1m, even as compared with cells expressing LMP1 DM, suggesting that unopposed TES2 signaling results in their downregulation (**Fig. 5B and S4A**). Cluster 3 genes were enriched for multiple KEGG metabolism pathways, including oxidative phosphorylation (**Fig. S5A**). Cluster 4 contained a smaller subset of host genes, downregulated by TES1 signaling, including in the WT LMP1 context. This gene set was enriched for SNARE interactions in vesicular transport and sphingolipid metabolism (**Fig. S5B**). Cluster 5 mRNAs were instead upregulated by unopposed TES2 signaling (**Fig. 5B, S5C**) and enriched for the KEGG ubiquitin mediated proteolysis pathway. Finally, Cluster 6 genes were repressed by TES1 and TES2 signaling in an additive manner (**Fig. 5D**). This gene set was enriched for mismatch and base excision repair, nucleotide metabolism, cystine and methionine metabolism. TES1 and TES2 signaling may additively recruit the same repressors or may instead recruit co-repressors to these sites. We also confirmed the concordance of our RNAseq through qRT-PCR of the transcript levels of CCL22, EBI3 and IRF4 in WT, TES1m, TES2m or DM induced expression (**Fig. S5 D-F**). As expected TES1m and TES2m significantly abrogate the expression of the genes while we barely detect anything for the DM.

We next used RNAseq to profile BL-41 Burkitt cells. RNAseq was performed at 24 hours post-expression of WT, TES1m, TES2m or DM LMP1. K-means analysis with n=6 clusters again revealed categories of genes that respond differently to LMP1 alleles (**Fig. S6A, Table S4-6**). As observed in Akata, Cluster 1 genes were most highly induced by WT, and to a lesser extent by TES1m or TES2m, suggesting that TES1/2 additively or synergistically induce their expression. Cluster 1 genes contained multiple pro-inflammatory factors, including chemokines and the interferon pathway transcription factors STAT1, IRF4, IRF5 and IRF9 (**Fig. S6A-B**). As observed in Akata, cluster 2 genes were induced more strongly by TES2m than by TES1m, and to a somewhat higher level by WT LMP1, suggesting that these are predominantly TES1 target genes (**Fig. S6C**). Consistent with our LCL and Akata cell analyses, CFLAR was a Cluster 2 gene more highly induced by LMP1 alleles with TES1 signaling, further underscoring it as a key TES1 target gene (**Fig. S6C**). By contrast, BL-41 Cluster 4 genes were instead suppressed by unopposed TES1 signaling even relative to levels observed in cells with LMP1 DM expression, suggesting that TES2 may block TES1 repressive effects on these host targets (**Fig. S6D**). Cluster 6 genes were enriched for the antigen presentation pathway and were most highly induced by TES2m, suggesting positive TES1 and potentially also negative TES2 roles in their induction (**Fig. S6E**). While concordant to a large degree, we speculate that observed differences between LMP1 effects on Akata versus BL-41 host gene expression may likely reflect the somewhat higher basal NF-κB levels observed in BL-41, and perhaps also differences in driver mutations pathways frequently found in EBV+ vs EBV-Burkitt lymphomas (69, 70). Nonetheless, both models highlight distinct clusters of host B-cell target genes that differ in responses to TES1, TES2 or combined TES1/2 signaling.

### LMP1 WT, TES1, TES2 and DM target genes

We next cross-compared the most highly differentially-expressed genes across the LMP1 conditions. At a fold-change >2 and adjusted p value <0.05 cutoff, WT LMP1 highly upregulated 1021 and downregulated 518 Akata genes, respectively. The most highly upregulated genes included multiple interferon stimulated genes, TRAF1, FAS and CFLAR (**Fig. 6A**). Interestingly, WT LMP1 decreased expression of the recombinase RAG1 and RAG2 mRNAs, as well as MME, which encodes CD10, a plasma membrane protein that we and others have found is downmodulated by EBV latency III (71-73). Enrichr analysis identified that EBV infection was the KEGG pathway most highly upregulated by WT LMP1 (**Fig. 6B**), reflecting the major LMP1 contribution to latency III datasets used in KEGG. Likewise, NF-κB and TLR signaling were also highly enriched, whereas downmodulated genes were enriched in the term primary immunodeficiency. Highly concordant effects were observed in the BL-41 cell context, where the same KEGG pathways were the most enriched amongst LMP1 upregulated genes (**Fig. S7A-C**). Cross-comparison of expression patterns in Akata and BL-41 with WT vs DM LMP1 expression again revealed highly concordant results (**Fig. 6C-D and S7D-F**), further validating a range of host genes as targets of TES1 and 2 signaling.

**Fig. 6.**
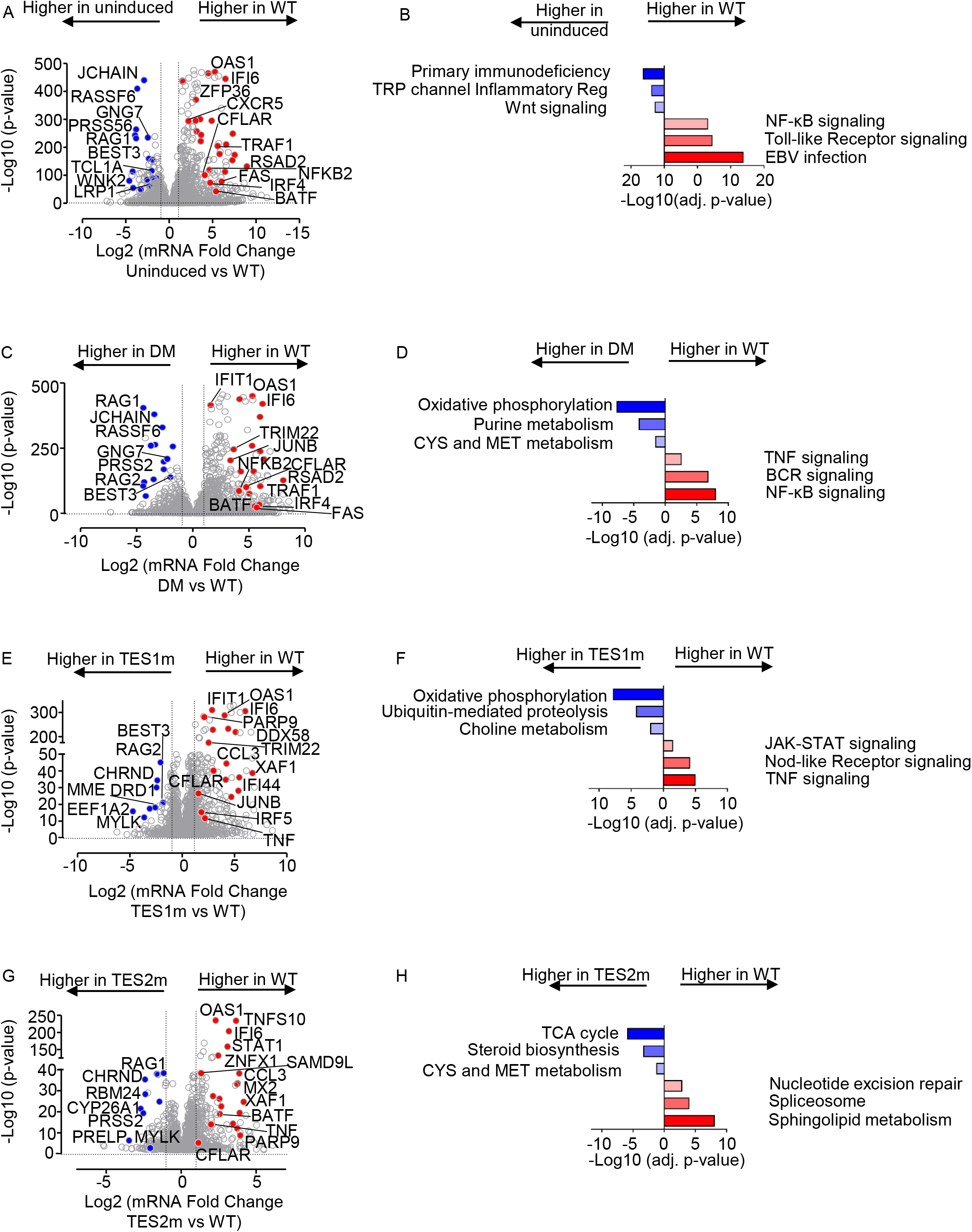
Characterization of Akata B-cell pathways targeted by TES1 vs TES2 signaling. (A) Volcano plot analysis of host transcriptome-wide genes differentially expressed in Akata cells conditionally induced for WT LMP1 expression for 24h by 250 ng/ml Dox versus in mock induced cells. Higher X-axis fold changes indicate genes more highly expressed in cells with WT LMP1 expression, whereas lower X-axis fold changes indicate higher expression in cells mock induced for LMP1. Data are from n=3 RNAseq datasets, as in **Fig. 5**. (B) Enrichr analysis of KEGG pathways most highly enriched in RNAseq data as in (A) amongst genes more highly expressed in Akata with WT LMP1 (red) vs amongst genes more highly expressed with mock LMP1 induction (blue). (C) Volcano plot analysis of host transcriptome-wide genes differentially expressed in Akata cells conditionally induced for WT vs DM LMP1 expression for 24h by 250 ng/ml Dox. Higher X-axis fold changes indicate genes more highly expressed in cells with WT LMP1 expression, whereas lower X-axis fold changes indicate higher expression in cells with DM LMP1. Data are from n=3 RNAseq datasets, as in **Fig. 5**. (D) Enrichr analysis of KEGG pathways most highly enriched in RNAseq data as in (C) amongst genes more highly expressed in Akata with WT LMP1 (red) vs amongst genes more highly expressed with DM LMP1 induction (blue). (E) Volcano plot analysis of host transcriptome-wide genes differentially expressed in Akata cells conditionally induced for WT vs TES1m LMP1 expression for 24h by 250 ng/ml Dox. Higher X-axis fold changes indicate genes more highly expressed in cells with WT LMP1 expression, whereas lower X-axis fold changes indicate higher expression in cells with TES1m LMP1. Data are from n=3 RNAseq datasets, as in **Fig. 5**. (F) Enrichr analysis of KEGG pathways most highly enriched in RNAseq data as in (E) amongst genes more highly expressed in Akata with WT LMP1 (red) vs amongst genes more highly expressed with TES1m LMP1 induction (blue). (G) Volcano plot analysis of host transcriptome-wide genes differentially expressed in Akata cells conditionally induced for WT vs TES2m LMP1 expression for 24h by 250 ng/ml Dox. Higher X-axis fold changes indicate genes more highly expressed in cells with WT LMP1 expression, whereas lower X-axis fold changes indicate higher expression in cells with TES2m LMP1. Data are from n=3 RNAseq datasets, as in **Fig. 5**. (H) Enrichr analysis of KEGG pathways most highly enriched in RNAseq data as in (G) amongst genes more highly expressed in Akata with WT LMP1 (red) vs amongst genes more highly expressed with TES2m LMP1 induction (blue).

To gain insights into how TES2 signaling shapes LMP1 genome-wide targets, we next cross-compared transcriptomes from Akata expressing WT vs TES1 mutant LMP1. At a fold-change >2 and adjusted p-value <0.05 cutoff, 561 genes were more highly expressed in WT than LMP1 TES1m, whereas 201 were less highly expressed. Interestingly, multiple interferon stimulated genes, including IFIT1, IFI6, STAT1 and IFI44 were amongst the most highly upregulated in WT LMP1 expressing cells (**Fig. 6E**). Enrichr analysis identified TNF, Nod-like receptor (NLR) and JAK/STAT signaling to be the most highly enriched KEGG pathways amongst genes more highly expressed in WT LMP1+ cells, whereas oxidative phosphorylation was the most highly enriched KEGG pathway amongst genes more highly expressed in cells expressing the TES1 mutant (**Fig. 6F**). Similar analyses on BL-41 cell datasets again revealed large numbers of differentially expressed genes in WT vs TES1m LMP1 expressing cells (**Fig. S8A-B** and **Table S5**).

To then gain insights into how TES1 signaling shapes LMP1 genome-wide target gene effects, we cross-compared Akata differentially expressed genes at 24 hours post expression of WT vs TES2 mutant LMP1. At a fold-change >2 and adjusted p value <0.05 cutoff, 275 genes were more highly expressed in Akata with WT than TES2 mutant LMP1, whereas 118 were less highly expressed. Once again, multiple interferon stimulated genes (ISG), including STAT1, IFI6 and OAS1 were more highly expressed in WT cells. Enrichr analysis identified sphingolipid signaling and metabolism to be most highly enriched KEGG pathways amongst genes upregulated genes, whereas TCA cycle was the most significant KEGG pathway amongst genes more highly expressed with TES2 mutant LMP1 expression (**Fig. 6G-H**). Similar numbers of genes were differentially regulated between WT and TES2m expressing cells in the BL-41 context, where Toll-like receptor signaling was the most highly enriched term amongst genes more highly expressed in WT LMP1+ cells (**Fig S8C-D** and **Table S6**). These analyses are consistent with a model in which TES1 and TES2 signaling additively or synergistically upregulate ISGs. Direct cross-comparison of TES1 vs TES2 signaling in Akata and BL41 further revealed pathways selectively targeted by either (**Fig. S8E-H**). In the Akata environment, cells expressing TES2m more highly induced ISGs, including IFIT1, IFI6, OAS, IFI44 and DDX58 (**Fig. S8E**). Enrichr analysis indicated that TES1 signaling most strongly induced the Nod-like receptor (NLR), necroptosis and chemokine signaling KEGG pathways. By contrast, TES2 signaling (from the TES1 mutant) most highly induced growth hormone and multiple amino acid metabolism KEGG pathways (**Fig. S8F**). In BL-41 cells, interferon stimulated genes were not as highly induced by TES2m (**Fig. S8G**). Since non-canonical NF-κB activity can strongly impact B cell type I interferon pathways (74), we suspect that differences in basal NF-κB activity in BL-41 may compensate to some extent to reduce this phenotype. Instead, cell adhesion molecules and TNF signaling were most highly enriched. For instance, CFLAR was significantly more highly induced by TES1 signaling, as was OTULIN, a deubiquitinating enzyme that controls TNF/NF-κB canonical pathway. FoxO and Toll-like receptor signaling were the most highly enriched KEGG pathways induced by TES2 signaling (by the TES1 mutant) in BL-41, with FOXO signaling the most selectively induced by TES1 mutant LMP1 (**Fig. S8H**).

We next directly cross-compared results from our LCL and Burkitt systems. Volcano plot analysis identified host cell genes whose expression was induced by Akata WT LMP1 expression but decreased by LCL LMP1 KO, suggesting that they are bone fide LMP1 targets (**Fig. S9A**, blue circles and **Table S1, S7**). This gene set included CFLAR, TRAF1, EBI3, CCL2, CD40, consistent with prior studies (27, 50, 55, 75-78). Similarly, genes whose expression was suppressed by Akata WT LMP1 expression but induced by LCL LMP1 KO were identified as LMP1-repressed host targets (**Fig. S9A**, red circles and **Table S1, S7**). We similarly cross-compared data from our Akata LMP1 expression and LCL LMP1 rescue datasets. Key targets of TES1 signaling, whose expression was significantly lower in Akata with TES1 mutant than WT LMP1 and also in LCLs rescued by TES1 mutant versus WT LMP1, included CFLAR, TRAF1, NFKB2 and CCL22 (**Fig. S9B** and **Table S2, S7**). Likewise, key TES2 targets more highly induced by WT than by TES2 mutant in both contexts included CCL22 and EBI3, whereas CR2, which encodes the EBV B-cell receptor complement receptor 2, was instead more highly expressed in cells with TES2m than WT LMP1 expression, suggesting it is repressed by TES2 signaling (**Fig. S9C** and **Table S3, S7**). Taken together, these findings serve to validate a class of host genes as LMP1 targets in the LCL context, although we cannot exclude that they are regulated through secondary effects.

### LMP1 TES1 and TES2 roles in LCL Dependency Factor BATF and IRF4 Expression

We next characterized LMP1 pathways important for BATF and IRF4 induction, given their key LCL but not Burkitt B-cell dependency factor roles (25, 79, 80). Notably, BATF and Jun family members bind cooperatively with IRF transcription factors to AP1-IRF composite DNA elements (AICE) (81), and JunB is the Jun family member predominantly expressed in LCLs (**Fig. 7A**). WT and TES2m LMP1 upregulated IRF4 mRNA abundance to a similar extent in Akata, whereas TES1m did so to a somewhat lesser extent. By contrast, TES1m and TES2m each upregulated BATF, but not quite as strongly as WT LMP1 (**Fig. 7B**). Taken together with the LCL LMP1 knockout data, these results suggest that LMP1 TES1 and TES2 signaling each support expression of the host transcription factors that bind to AICE.

**Fig. 7.**
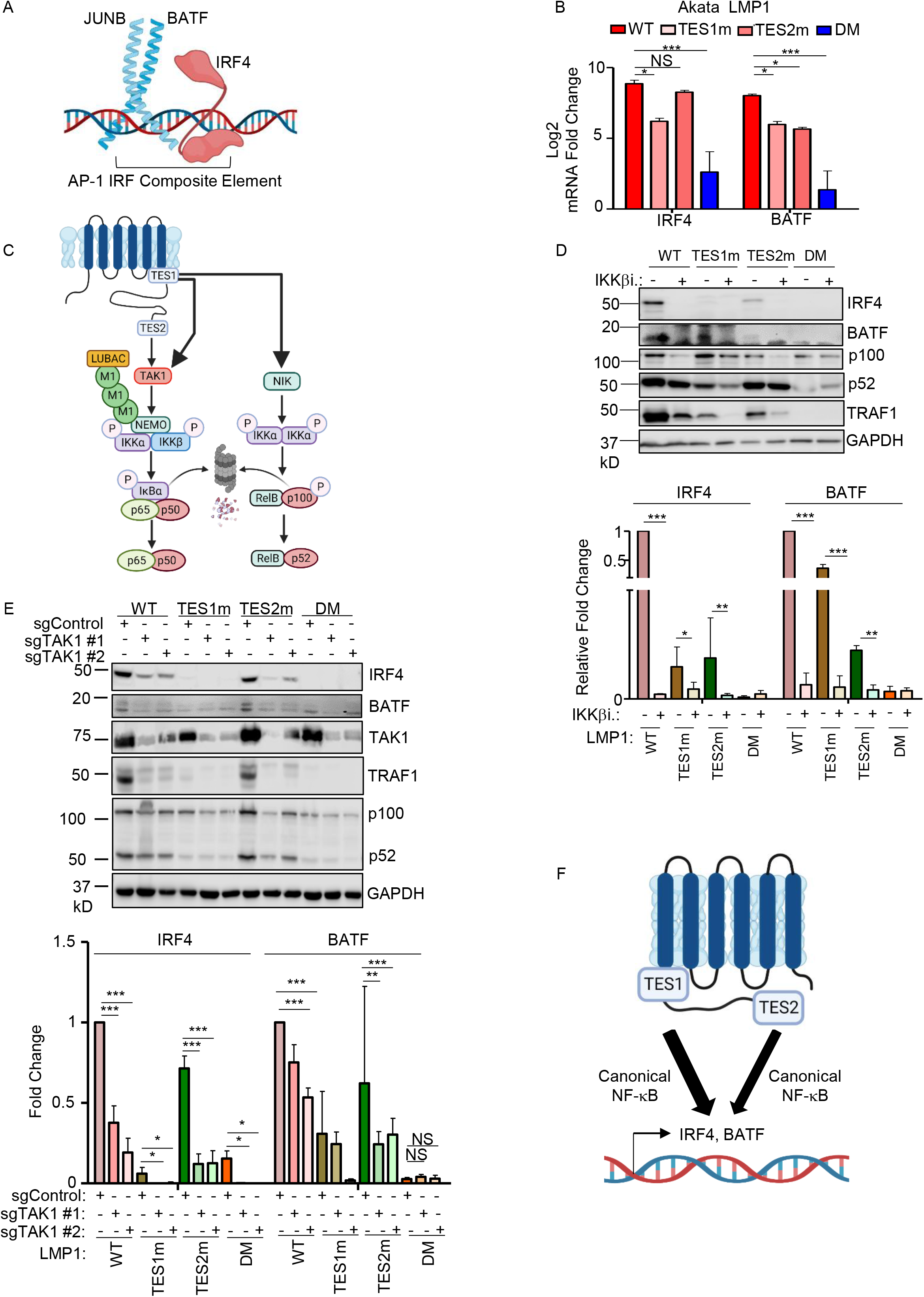
Roles of TES1 and TES2 canonical NF-κB pathways in LCL dependency factor BATF and IRF4 expression. (A) Schematic diagram of JUNB, BATF and IRF4 at an AP-1/IRF composite DNA site. (B) Mean + SD fold changes of IRF4, BATF and JUNB mRNA abundances from n=3 RNAseq replicates of Akata cells expressing the indicated LMP1 cDNA for 24 hours, as in Fig. 5. p-values were determined by one-sided Fisher’s exact test. *p<0.05, ***p<0.001. (C) Schematic diagram of LMP1 TES1 and TES2 NF-κB pathways. TES1 and TES2 each activate canonical NF-κB pathways, whereas TES1 also activates non-canonical NF-κB. (D) Immunoblot analysis of WCL from Akata cells induced for LMP1 expression by 250 ng/ml Dox for 24 hours, either without or with 1 μM IKKβ inhibitor VIII. Shown below are relative fold changes + SD from n=3 replicates of IRF4 or BATF vs GAPDH load control densitometry values. Values in vehicle control treated WT LMP1 expressing cells were set to 1. P-values were determined by one-sided Fisher’s exact test. **p<0.001, ***p<0.0001. (E) Immunoblot analysis of WCL from Cas9+ Akata cells expressing control or either of two *TAK1* targeting sgRNAs, induced for LMP1 expression by 250 ng/ml Dox for 24 hours. Shown below are relative foldchanges + SD from n=3 replicates of IRF4 or BATF vs GAPDH load control densitometry values. Levels in cells with control sgRNA (sgControl) and WT LMP1 were set to 1. **p<0.001, ***p<0.0001. (F) Model of additive TES1 and TES2 canonical NF-κB pathway effects on BATF and IRF4 induction.

We next examined the contribution of LMP1 NF-κB pathways to B cell IRF4 and BATF expression. LMP1 CRISPR knockout in GM12878 LCLs or in latency III Jijoye Burkitt cells significantly reduced BATF and to IRF4 expression at the protein level, though effects on IRF4 were more subtle at this early timepoint (**Fig. S10A**). LMP1 induction of IRF4 and BATF was strongly impaired by induction of either TES1m or TES2m in Akata or in LCLs, relative to levels in cells with WT LMP1 (**Fig. 7B and S10B**). To test canonical NF-κB pathway roles in IRF4 and BATF induction, we then induced LMP1 in the absence or presence of a small molecule antagonist of the kinase IKKβ, which is critical for canonical NF-κB pathway signaling. IKKβ inhibition blocked their residual induction by TES1m and TES2m (**Fig. 7C-D**). The IKKβ inhibitor also reduced BATF and IRF4 expression in GM12878 (**Fig. S10C**). Similar results were obtained in Akata cells that co-induced LMP1 with an IκBα super-repressor (IκBα-SR), in which IκBα serine 32 and 36 to alanine point mutations prevent its canonical pathway phosphorylation and proteasomal degradation (**Fig. S10D**). Furthermore, CRISPR KO of the canonical NF-κB pathway kinase TAK1 significantly impaired IRF4 induction by WT and also TES2m LMP1 (**Fig. 7E**). Taken together, these results suggests that canonical NF-κB pathways driven by both TES1 and TES2 signaling are each important for BATF and IRF4 expression (**Fig. 7F**).

### LMP1 TES1 and TES2 roles in EBV Super-Enhancer target induction

The five LMP1-activated NF-κB transcription factor subunits and four EBNAs target a set of LCL host genome enhancers termed EBV super-enhancers (EBV SE). EBV SE are characterized by occupancy by all five NF-κB subunits, EBNAs 2, LP, 3A and 3C and markedly higher and broader histone H3K27ac ChIP-seq signals than at typical LCL enhancers (**Fig. 8A**). SE are critical for cell identity and oncogenic states (82), and EBV SE are important for LCL growth and survival (83-85). However, little has remained known about the extent to which LMP1 TES1 versus TES2 signaling contribute to EBV SE. To gain insights, we therefore interrogated EBV SE gene target responses, we first plotted EBV SE gene target responses to GM12878 LMP1 CRISPR KO. At the early timepoint of 48 hours after LMP1 editing, expression of SE targets TRAF1 is significantly decreased while expression of PRDM1 (which encodes the transcription repressor BLIMP1) and GPR15 (which encodes a G-protein coupled chemokine receptor) each is significantly increased, though most other EBV SE gene targets did not significantly change at this early timepoint (**Fig. 8B**, red circles and **Table S7**). This data suggests that EBV SE are robust to short-term perturbations of LMP1 expression, perhaps given persistence of the established epigenetic landscape built at these key sites.

**Fig. 8.**
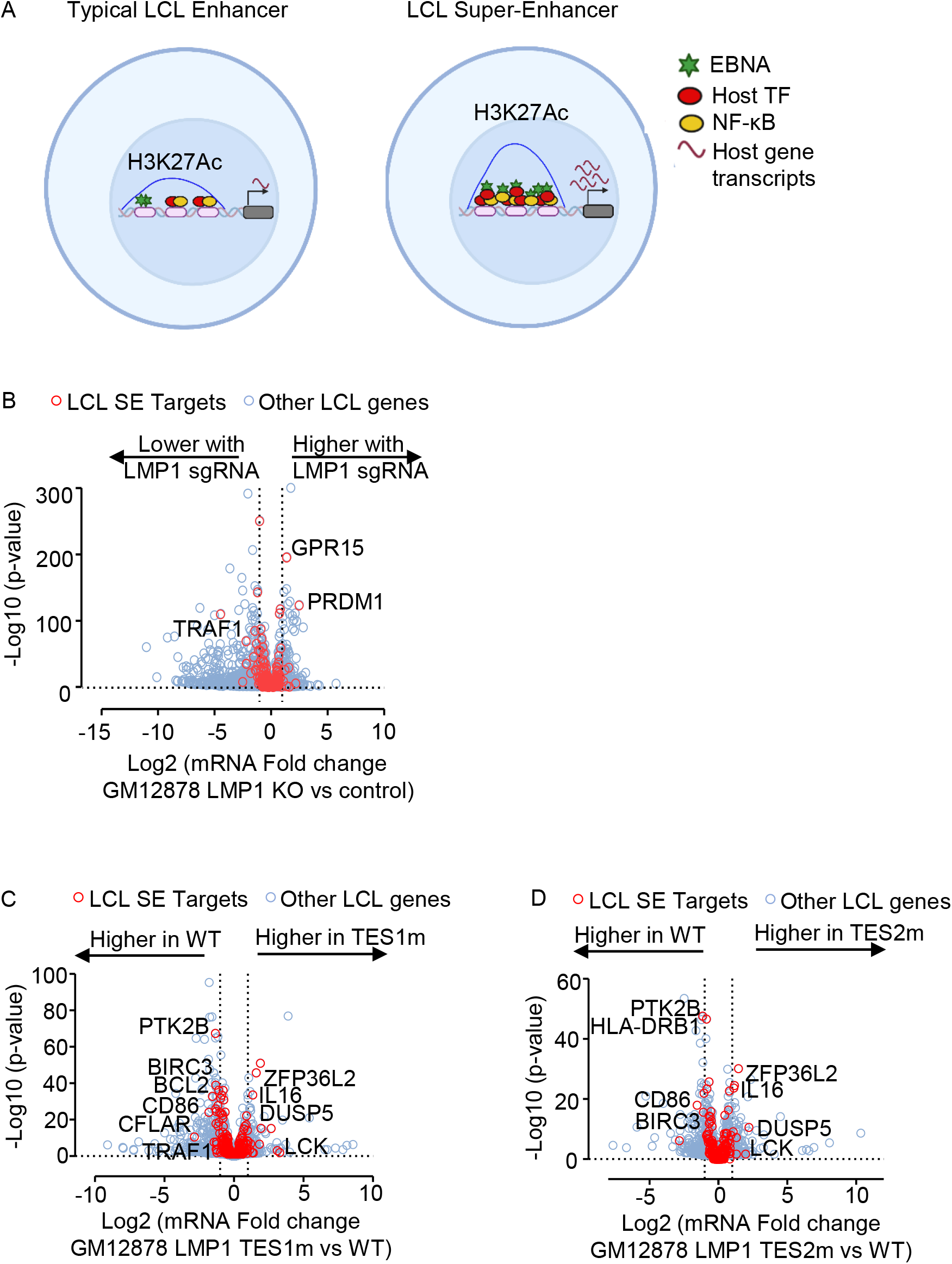
LMP1 TES1 and TES2 roles in EBV super-enhancer target gene regulation in GM12878 LCLs. (A) Schematic diagram of typical LCL enhancers vs super-enhancers. Super-enhancers have significantly broader and taller histone 3 lysine 27 acetyl (H3K27Ac) peaks. EBV SE are host genomic enhancer sites bound by all five LMP1-activated NF-κB transcription factor subunits, EBNA-2, LP, 3A, and 3C. (B) Volcano plot analysis of mRNA values in GM12878 expressing LMP1 vs control sgRNAs as in **Fig. 1**. Genes targeted by EBV super-enhancers (SE) are highlighted by red circles, whereas other LCL genes are indicated by blue circles. Genes more highly expressed with LMP1 KO have higher x-axis values, whereas those downmodulated by LMP1 KO have lower values. P value <0.05 and >2-fold gene expression cutoffs were used. (C) Volcano plot analysis of mRNA values in GM12878 expressing LMP1 sgRNA with TES1m versus WT LMP1 cDNA rescue, as in **Fig. 2-3**. Genes targeted by EBV super-enhancers (SE) are highlighted by red circles, whereas other LCL genes are indicated by blue circles. Genes more highly expressed with endogenous LMP1 KO and TES1m rescue have higher x-axis values, whereas those more highly expressed with WT LMP1 rescue have lower values. P value <0.05 and >2-fold gene expression cutoffs were used. (D) Volcano plot analysis of mRNA values in GM12878 expressing LMP1 sgRNA with TES2m versus WT LMP1 cDNA rescue, as in (C). Genes targeted by EBV super-enhancers (SE) are highlighted by red circles, whereas other LCL genes are indicated by blue circles. Genes more highly expressed with endogenous LMP1 KO and TES2m rescue have higher x-axis values, whereas those more highly expressed with WT LMP1 rescue have lower values. P value <0.05 and >2-fold gene expression cutoffs were used.

To then identify the extent to which LMP1 is sufficient to alter expression of these LCL EBV SE targets, we next visualized effects of WT, TES1m or TES2m rescue of LMP1 KO LCLs. Potentially because this analysis could be done at a later timepoint post LMP1 CRISPR editing (day 6 post LMP1 KO), we observed stronger effects on EBV SE target gene expression. Rescue with TES1 mutant LMP1 caused significant loss of CFLAR, BCL2 and TRAF1 relative to levels with WT rescue, whereas rescue with either TES mutant significantly lowered levels of CD86 and BIRC3 messages from levels observed with WT rescue (**Fig. 8C-D** and **Table S8-9**). As a complementary approach, we also analyzed effects of conditional LMP1 induction in our Burkitt models on EBV SE target gene expression. In Akata B cells, WT LMP1 induction was sufficient to significantly upregulate the abundance of the majority of EBV SE target gene mRNAs (**Fig. S11A** and **Table S1**). This result suggests that while EBNA-2, LP, 3A and 3C also target these sites in LCLs, LMP1 can independently alter expression of most EBV SE targets, albeit not necessarily to the same extent as latency III. By contrast, Akata TES1m or TES2m induction less strongly induced most EBV SE targets (**Fig. S11B-C**). A similar pattern was observed in BL-41 cells (**Fig. S11D-F** and **Table S4**). Taken together, our results indicate that TES1 and TES2 likely play key joint roles in the induction of EBV SE target genes.

## Discussion

Why the LMP1 C-terminal tail TES1 and TES2 domains are each necessary for lymphoblastoid B-cell immortalization, and whether each are necessary for LCL survival have remained longstanding questions. To gain insights into key LMP1 B-cell roles, we used a novel LCL LMP1 KO with conditional LMP1 rescue system to identify that signaling by TES1, but not TES2, is required for LCL survival. We performed systematic B-cell transcriptome-wide analyses to identify effects of LMP1 knockout in the absence or presence of rescue by WT, TES1 mutant or TES2 mutant LMP1, at early timepoints where cells remained viable. These highlighted key LMP1 TES1 and TES2 roles in support of LCL dependency factor and EBV super-enhancer target gene expression. As a complimentary approach to identify host B-cell genome-wide targets of LMP1 signaling, we also profiled EBV-negative Burkitt B-cell responses to conditional expression of wildtype LMP1, or LMP1 point mutants abrogated for signaling by TES1 or TES2. As has previously been described in comparisons of LMP1 vs LMP2A expression(86), TES1 and TES2 signaling effects were not simply additive, but yielded distinct effects on host target genes, with either TES domain more strongly inducing or repressing genes at particular sites, but opposing one another at other sites to fine tune target gene expression (**Fig. 9**).

**Fig. 9.**
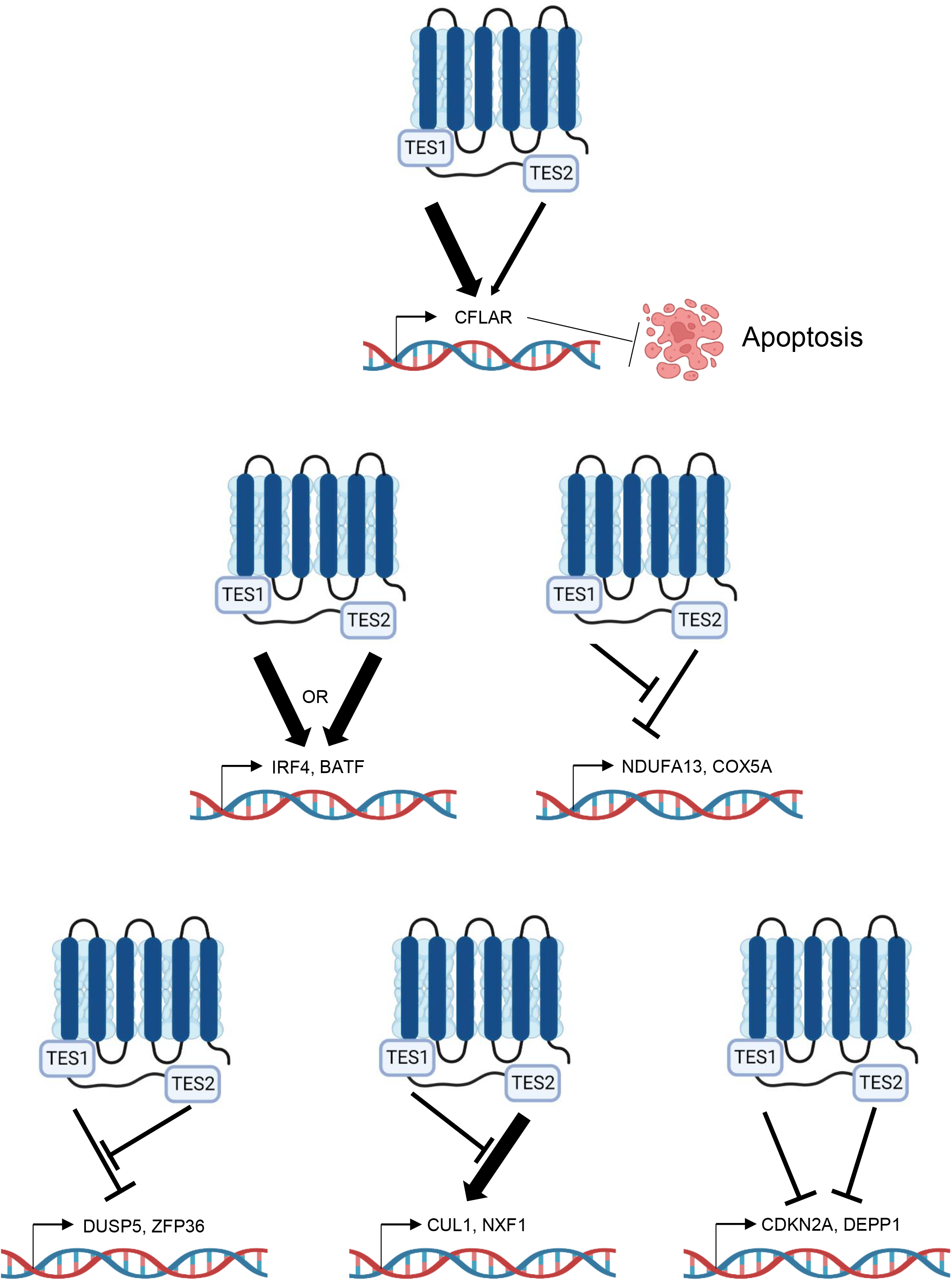
Model highlighting different modes of LMP1 TES1 and TES2 cross-talk in B-cell target gene regulation.

LMP1 KO rapidly altered expression of ∼3400 LCL genes, with roughly similar numbers of host genes being downregulated as upregulated. This result is consistent with prior microarray analyses of LMP1 targets in 293 cells induced for TES2 signaling and in Burkitt-cells induced for LMP1 (27, 50), as well as in microarray analysis of EBV-infected B-cells at timepoints where LMP1 expression increases(87), further suggesting that LMP1 strongly remodels the transcriptome by pleotropic effects on host gene expression. Similar numbers of LMP1 targets were also found in microarray profiling of B-cells with transgenic LMP1 expression (77). While TES1 and TES2 induce host genes through activation of NF-κB, MAP kinase, PI3K and interferon regulatory factor pathways (3-5, 11-20), comparatively little is known about how LMP1 downmodulates target gene expression. However, one mechanism by which LMP1 may repress target genes could be through NF-κB complexes, including p50 or p52 homodimers, or p50:52 heterodimers, potentially together with BCL3 (88, 89), as these NF-κB complexes lack transactivation domains. We do not suspect that these changes were secondary to cell death, as we performed profiling on viable cells at an early timepoint post-CRISPR editing. However, the result that LMP1 is critical for LCL survival builds on prior analyses, which showed that blockade of LMP1/NF-κB signaling triggers LCL apoptosis (50, 87, 90, 91).

Conditional expression of WT LMP1 rescued LCL survival, confirming on-target CRISPR effects on EBV genomic LMP1. Our rescue approach identified that loss of TES1, but not TES2 signaling, triggered LCL apoptosis, as judged by upregulation of caspase 3 and 7 activity and by FACS analysis for plasma membrane Annexin V. Disruption of cell death signaling is a hallmark of cancer (92), and enrichment analysis identified that the KEGG apoptosis pathway was highly altered by loss of TES1. Notably, LMP1 has thus far remained an undruggable target. Therefore, these results suggest that small molecule or peptide inhibitors that block TES1 signaling may have therapeutic benefit, even in the absence of effects on TES2, for instance in the setting of EBV-driven post-transplant and central nervous system lymphomas, which frequently express the latency III program and which are modeled by LCLs. It will be of interest to determine whether TES1 signaling has similarly important roles in apoptosis blockade in other EBV-infected tumor contexts, including in Hodgkin lymphoma Reed-Sternberg tumor cells and in nasopharyngeal carcinoma, where little is presently known about TES1 vs TES2 roles. The LCL dependency factor CFLAR, which encodes the extrinsic apoptosis pathway inhibitor c-FLIP, was highly downmodulated upon loss of TES1 signaling, to a significantly greater extent than upon loss of TES2 signaling. This is consistent with prior microarray analysis identified that identified cFLIP as an LMP1 target (50), which we now mostly induced by TES1 signaling. We previously identified that c-FLIP is required for LCL survival and is required to block the extrinsic apoptosis pathway that is otherwise triggered by TNFα signaling, likely in response to EBV oncogenic stress (25). Therefore, our data newly identified CFLAR as a key TES1 target gene and suggest that TES1 signaling is required for LCL survival, at least in part due to obligatory roles in cFLIP induction. Our data raises the interesting question of why CFLAR expression is particularly dependent on TES1 signaling, to a much greater extent than TES2. It is plausible that a TES1-driven non-canonical NF-κB pathway is particularly important for cFLIP transcription. However, TES1 signaling also strongly activates canonical NF-κB pathways (13, 15, 89), which may be critical for CFLAR induction. Alternatively, MAP kinases or PI3K activated by TES1 (4, 36, 79, 88, 93-95) may also support CFLAR expression.

LMP1 expression both activates and blocks apoptosis (42, 43, 96), and our data suggests that TES1 induction of CFLAR is central to this balance, perhaps together with BCL2 family members such as BFL1 (96, 97). However, in addition to targeting CFLAR, apoptosis pathways were enriched amongst LMP1 target genes. It has also been reported that the six LMP1 transmembrane domains induce apoptosis through activation of an unfolded protein response, while LMP1 C-terminal domain signaling counteracts this (96). Similarly, LMP1 induces *c-jun*, *junB* and *junD* (98), which may play roles in balancing proliferation and apoptosis responses (99). LMP1 also closely regulates the expression of pro-apoptotic and anti-apoptotic genes to allow for cell proliferation (98). We also observed downregulation of the p53 antagonist MDM2 and upregulation of p53 upon LMP1 KO. Both TES1 and TES2 had important roles in regulation of MDM2 expression. Thus, taken together with prior studies, our data sheds light into the balance of cell death and survival signals triggered by LMP1 signaling.

Our data further highlight LMP1 TES1 and TES2 roles in support of additional LCL dependency factors, in particular BATF and IRF4. In contrast to CFLAR, TES1 and TES2 signaling were each found to be important for BATF and IRF4 expression, both in LCL and Burkitt cell models. TES1 and TES2 driven canonical NF-κB signaling supported BATF induction in EBV-negative Burkitt cells, where EBNA2 is not expressed. BATF and IRF4 expression rapidly decreased upon LMP1 KO in LCLs, even upon rescue by LMP1 signaling from only one TES domain. Thus, in LCLs, EBNA2 and LMP1-stimulated canonical NF-κB jointly induce this critical LCL dependency factor. LMP1 canonical NF-κB pathways were also critical for inducing IRF4, which binds with BATF to composite AICE DNA sites. As EBNA2 also supports BATF (100) and EBNA3C also supports IRF4 expression (101), our results further highlight BATF and IRF4 as major hubs of EBV oncoprotein cross-talk.

We previously used ChIP-seq to characterize the LCL NF-κB genomic binding landscape (32). Rather than identifying readily recognizable LMP1 canonical versus non-canonical NF-κB target genes, this study identified complex patterns of occupancy by the five NF-κB transcription factor subunits at LCL enhancers and promoters. However, LCL enhancers often target multiple genes, often from long distances (83), complicating cross-comparison with this study. Furthermore, concurrent LMP1 TES1 and TES2 signaling yields up to 13 distinct NF-κB transcription factor dimers in LCL nuclei (32), including dimers such as cRel:p52 that are under control of both NF-κB pathways. Conditional expression of TES1m or TES2m should yield a considerably less complex NF-κB. Therefore, a future objective will therefore be to perform NF-κB ChIP-seq in using the conditional TES1m and TES2m conditional Burkitt models reported here, as these may yield less complex patterns of NF-κB occupancy.

Our analyses highlight independent, shared and antagonistic LMP1 TES1 and TES2 roles in B-cell genome-wide target gene regulation (**Fig. 9**). How independent or combined TES1 and TES2 signaling have different effects on clusters of target genes will be important to define. Our results suggest multiple testable models. For instance, with regards to genes which were induced weakly by TES2 signaling, somewhat more by TES1 signaling but more highly by WT LMP1, we speculate that TES1 and TES2 may cross-talk at the epigenetic level. For example, our results are consistent with a model in which TES1 signaling increases chromatin accessibility at these sites, including the genes encoding IFIT1 and CXCL9. Once accessible, both TES1 and TES2 signal-dependent pathways may then additively or perhaps synergistically upregulate these sites. Alternatively, TES1 signaling could be needed to dismiss a repressor, such that these sites can then be stimulated by both LMP1 domains. TES1 signaling may also activate a key positive regulator such as BCL3 (88), which then functions together with transcription factors activated by TES1 and TES2 pathways. By contrast, a large number of genes appeared to be additively repressed by TES1 and TES2 signaling. TES1 and TES2 may recruit co-repressors to these sites, may additively recruit the same repressor or may reduce chromatin accessibility.

We also identified a cluster of LMP1 response genes, in which unopposed TES2 signaling caused target gene downregulation (**Fig. 9**). In Akata cells, this cluster (Cluster 3) was enriched for metabolism genes, raising the possibility that another key TES1 signaling role is to support metabolic pathway remodeling by EBV, such as glutathione metabolism or OXPHOS (86, 102-104). It is possible that TES2 induces a repressor that targets these sites, but that TES1 signaling serves to blunt its induction. Alternatively, TES1 signaling may induce an activator that counter-balances TES2-driven repressor activity. Or, TES2 signaling may reduce chromatin accessibility at these sites in the absence of TES1. By contrast, genes in Akata Cluster 5 were upregulated by unopposed TES2 signaling, but not by WT LMP1 or unopposed TES1 signaling (**Fig. 9**). TES1 signaling may instead recruit repressors or alter chromatin accessibility at these sites. Epigenetic analyses of histone repressive marks, such as ChIP studies of H3K9me3 and H3K27me3, as well as ATAC-seq studies of DNA packaging, should help to differentiate between these and other possibilities.

Our studies provide insights into TES1 and TES2 roles in regulation of B-cell EBV SE targets. Although EBV SE are highly co-occupied by all five LMP1-activated NF-κB subunits, individual TES1 and TES2 roles in EBV SE target gene regulation has remained unstudied. Interestingly, while either TES1 or TES2 signaling was sufficient to induce many EBV SE targets, TES1 induced a larger number, perhaps because it highly induces both canonical and non-canonical pathways and therefore activates all 5 NF-κB subunits. By contrast, LMP1 KO perturbed expression of only a small number of EBV SE targets in LCLs, likely because of the early timepoint profiled prior to cell death, which left little time for epigenetic remodeling of these sites.

WT or DM LMP1 expression caused highly concordant changes in host gene expression in Akata and BL-41 Burkitt models. We speculate that differences in response to signaling by TES1 or TES2 alone may have instead arisen from distinct host genome mutation landscapes between these two human tumor derived models, which alter the basal NF-κB level.

Nonetheless, since EBV can infect a wide range of B-cells, including of distinct differentiation or activation states that alter NF-κB states, differences between Akata and BL-41 provide insights into how LMP1 may function in differing human B-cell contexts, and suggest that LMP1 may have evolved signaling by both TES domains to increase robustness across the spectrum of infected B-cell states.

LMP1 polymorphisms have been observed across EBV strains, in particular between type I and II EBV (105). In addition, LMP1 C-terminal tail polymorphisms have been observed in several analyses of EBV genomes isolated from Hodgkin lymphoma and nasopharyngeal carcinoma tumor cells, though the roles of these variant LMP1 sequences remain controversial. Amongst the best studied is a 30 base pair deletion present in the EBV CAO and 1510 strains isolated from Asian NPC tumors, which causes loss of LMP1 residues 343-352 (106, 107). This 30bp deletion has also been reported as enriched in EBV genomes isolated from Hodgkin tumor samples (107-110). A meta-analysis of 31 observational studies suggested a possible association between this LMP1 C-terminal tail deletion and nasopharyngeal carcinoma susceptibility, but was limited by small sample size and considerable variation between studies (111). Deletion of these residues, which comprise the last eight residues between the TES1/CTAR1 and the first two residues of TES2/CTAR2, enhance rodent fibroblast transformation by LMP1 (112) and may reduce immunogenicity (113), but were not found to enhance LMP1-mediated NF-κB activation (114). These strains also have multiple additional LMP1 amino acid polymorphisms, which are instead implicated in enhanced NF-κB activation, and were mapped to the LMP1 transmembrane domains (114). Little information is presently available about how these polymorphisms alter LMP1 target gene expression. It will therefore be of interest to use the approaches presented here to characterize how LMP1 polymorphisms present in tumor-derived EBV strains may alter transcriptome responses to TES1 and TES2 signaling.

In summary, we identified LMP1 genome-wide B-cell targets and characterized their responses to signaling by TES1 and/or TES2. Signaling by TES1, but not TES2 was identified to be critical for blockade of LCL apoptosis, and CFLAR was identified as the LCL dependency factor most strongly impacted by shutoff of TES1 signaling as opposed to TES2. CRISPR KO approaches highlighted LCL genes that are highly sensitive to perturbation of TES1 and/or TES2 signaling. K-means analysis highlighted gene clusters with distinct expression responses to signaling by one or both LMP1 transformation essential domains in the latency III LCL versus EBV-negative Burkitt B-cell contexts. These studies highlight multiple levels by which TES1 and TES2 signaling alter LMP1 target gene expression, including by additive vs opposing roles. Collectively, these studies provide new insights into key non-redundant versus joint TES1 and TES2 roles in B-cell target gene regulation and highlight TES1 signaling as a key lymphoblastoid B-cell therapeutic target.

## Supporting information

Supplemental Table 1_ Akata Induced LMP1 WT vs mock Induced

Supplemental Table 2_ Akata Induced LMP1 TES1m vs Induced WT

Supplemental Table 3_ Akata Induced LMP1 TES2m vs Induced WT

Supplemental Table 4_BL-41 Induced LMP1 WT vs mock Induced

Supplemental Table 5_BL-41 Induced LMP1 TES1m vs Induced WT

Supplemental Table 6_ BL-41 Induced LMP1 TES2m vs Induced WT

Supplemental Table 7_ LCL LMP1 KO vs Control

Supplemental Table 8_ Induced LMP1 TES1m versus Induced WT

Supplemental Table 9_ Induced LMP1 TES2m versus Induced WT

## Acknowledgements

This work was supported by NIH R01 CA228700 and AI164709, U01 CA275301, P01 CA269043 and a Burroughs Wellcome Career Award in Medical Sciences to B.E.G, T32 T32AI007245 and an American Cancer Society Post-doctoral Fellowship to E.M.B, T32AI007061 to N.B., T32 T32AI007245 and F32 AI172329 to L.A.M.N and NIH K99 1K99DE031016 to R.G. All RNAseq and proteomics datasets will be published in the GEO omnibus GSE228158, GSE228167, GSE228178 and GSE240732 upon manuscript publication.

## AUTHOR CONTRIBUTIONS

B.M and E.M.B performed the experiments. B.M, N.R.B., R.G, E.M.B and L.A.M.N performed bioinformatics analyses. B.M and B.E.G designed the experiments. B.M and B.E.G wrote the manuscript. All authors analyzed the results, read and approved the manuscript.

## Materials and Methods

### Cell lines, culture, and vectors

HEK293T cells were purchased from ATCC and cultured in Dulbecco’s modified Eagle medium (DMEM, Gibco) with 10% fetal bovine serum (FBS, Gibco). EBV-negative Akata and BL-41 cells were obtained from Elliott Kieff; GM12878 were purchased from Coriell. Mutu I was obtained from Jeff Sample and Jijoye was purchased from ATCC. All B-cell lines stably expressed *Streptococcus pyogenes* Cas9 and were grown in Roswell Park Memorial Institute (RPMI) 1640 (Life Technologies) with 10% fetal bovine serum (Gibco) and penicillin-streptomycin in a humidified chamber with 5% carbon dioxide. LMP1 wildtype, TES1 alanine point mutant 204PQQAT208 -> AQQAT, TES2 384YYD386->ID mutant and double mutant LMP1 with both AQQAT and ID mutations were cloned into the pLIX-402 vector. pLIX-402 uses a TET-On TRE promoter to drive transgene expression and a C-terminal HA-tag fusion. Lentivirus vectors were used to establish stable Cas9+/GM12878, Cas9+/EBV-Akata and Cas9+/EBV-BL-41 Burkitt cells. Cell lines were then maintained with 0.5 μg/mL puromycin or 25 μg/mL hygromycin. For LMP1 inducible expression studies, 0.5X10^6 cells/ml were plated on Day-1 in 2ml of fresh RPMI in a 12-well plate. Cell were treated with 250ng/ml (in Burkitt cell models) for 24 hours prior to sample collection for downstream analyses or with 400ng/ml doxycycline (in LCL LMP1 cDNA rescue model) (Sigma #D9891) to allow for LMP1 rescue upon CRISPR knockout of LMP1. The Iκβα Super-repressor (SR) lacking residues 1-67 has previously been reported (27).

### Antibodies and Reagents

Cell Signaling Technology (CST) TRAF1 (#4715, Rabbit mAb), p105/50 (#3035, Rabbit mAb), RelA (#8242, Rabbit mAb), phospho-RelA (Ser536) (3033, Rabbit mAb), RelB (#4922, Rabbit mAb), cRel (#4727, Rabbit mAb), IκBα (#9247, Mouse mAb), IRF4 (#4964, Rabbit mAb), TAK1 (#4505, Rabbit mAb), V5 (#13202, Rabbit mAb), HRP-linked anti-mouse IgG(7076), FLIP (#8510, Rabbit mAb), HRP-linked anti-rabbit IgG (#7074) were used in this study at 1:1000 dilution. p100/52 (EMD Millipore #05-361, Mouse mAb, 1:1000), GAPDH (EMD Millipore #MAB374, Mouse mAb, 1:500) was used. S12 mouse monoclonal antibody against LMP1 was purified from hybridoma supernatant (115). The IKKβ inhibitor IKK-2 inhibitor VIII (ApexBio, #A3485), puromycin dihydrochloride (Thermo Fisher #A1113803), and hygromycin B (Millipore #400052).

### Growth Curve Analysis

For growth curve analysis, cells were counted and then normalized to the same starting concentration, using the CellTiterGlo (CTG) luciferase assay (Promega, Cat#G7570). Live cell numbers were quantitated at each timepoint by CTG measurements, and values were corrected for tissue culture passage. Fold change of live cell number at each timepoint was calculated as a ratio of the value divided by the input value. For the Caspase-Glo 3/6 Assay (Promega #G8092), Caspase-Glo 3/7 reagent was added to the cells, mixed and incubated for 30 minutes followed by recording of the luminescence. Readings were normalized to respective CTG values of the samples that were performed and collected concurrently.

### CRISPR/Cas9 editing

B-cell lines with stable Cas9 expression were established as described previously (47). Briefly, HEK293T cells were plated at a density of 300,000 cells per well in 2 mL DMEM supplemented with 10% FBS on Day -1. The following day (Day 0) plated cells were transfected with the TransIT-LT1 Transfection Reagent (Mirus #2306), according to the manufacturer’s protocol. Transfection media was replaced by RPMI 16 hours later (Day 1). B-cells were plated at 1.2X 10^6 density in a 6-well plate on Day 1. Lentivirus collected on Day 2 were added to the B-cells for spinoculation at 2000rpm for 2 hours at 37C and 4μg/ml of polybrene. Spinoculated cells were placed in in a humidified chamber with 5% carbon dioxide for 6 hours, then pelleted and resuspended in fresh RPMI/FBS. 48 hours post-transduction, transduced cells were selected by addition of puromycin 3 μg/ml or 200 μg/ml hygromycin. Broad Institute pXPR-515 control sgRNA (targets a non-coding intergenic region), Avana or Brunello library sgRNAs, as listed in Table 1, were cloned into lentiGuide-Puro (Addgene, catalog #52963) or pLenti SpBsmBI sgRNA Hygro (Addgene, catalog #62205).

**Table 1.**
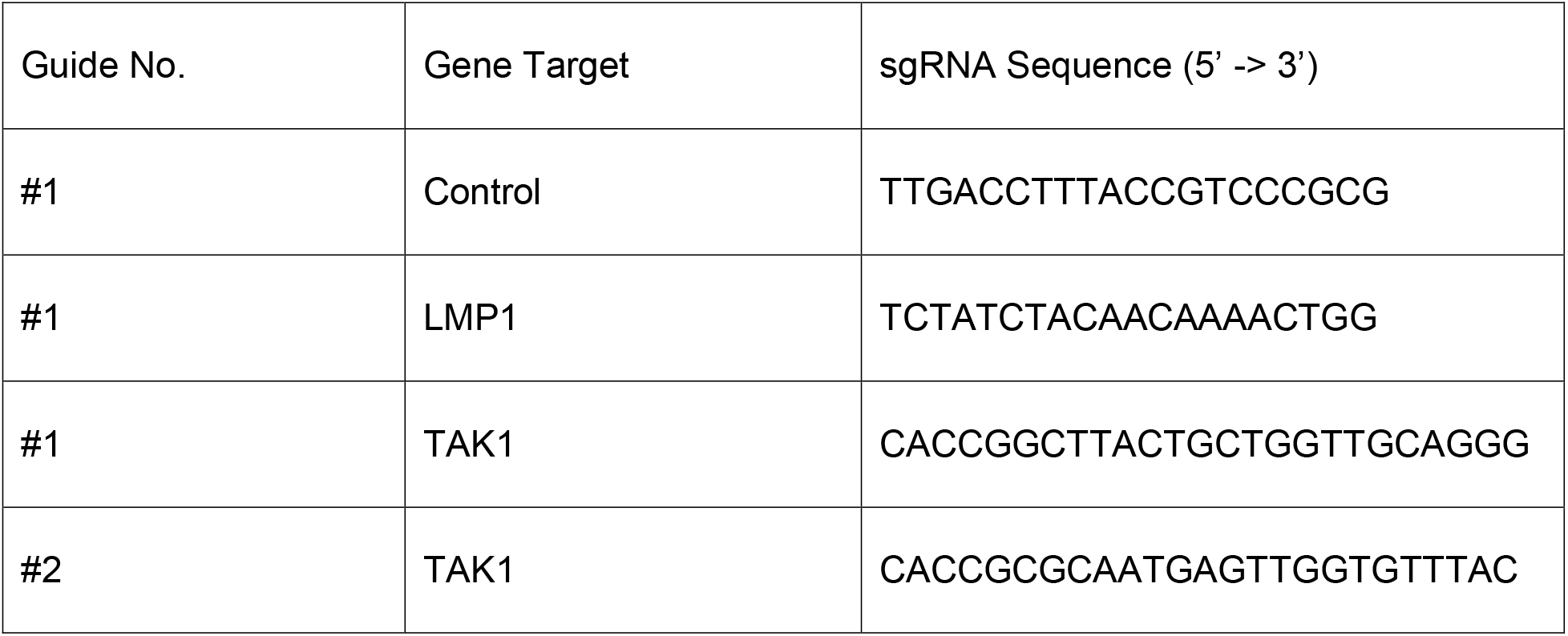
sgRNAs used in this study.

### RNAseq

Total RNA was isolated using RNeasy mini kit (Qiagen #74106) with in-column genomic DNA digestion step (RNAse-free DAase set, Qiagen #79254) according to the manufacturer’s protocol. To construct indexed libraries, 1 μg of total RNA was used for polyA mRNA selection using NEBNext Poly(A) mRNA Magnetic Isolation Module (Cat#E7490S), and library preparation with NEBNext Ultra RNA Library Prep with Sample Purification Beads (Cat#E7765S). Each experimental treatment was performed in biological triplicate. Libraries were multi-indexed (NEB 7335L and E7500S), pooled and sequenced on an Illumina NextSeq 500 sequencer using single 75 bp read length. Adaptor-trimmed Illumina reads for each individual library were mapped back to the human GRCh37.83 transcriptome assembly using STAR2.5.2b (116). FeatureCounts was used to estimate the number of reads mapped to each contig (117). Only transcripts with at least 5 cumulative mapping counts were used in this analysis. DESeq2 was used to evaluate differential expression (DE) (118). DESeq2 uses a negative binomial distribution to account for overdispersion in transcriptome data sets. It is conservative and uses a heuristic approach to detect outliers while avoiding false positives. Each DE analysis was composed of a pairwise comparison between experimental group and the control group. Differentially expressed genes were identified after a correction for false discovery rate (FDR). For more stringent analyses, we set the cutoff for truly differentially expressed genes as adjusted p value (FDR corrected) < 0.05 and absolute fold change > 1.5. DE genes meeting this cutoff were selected and subject to downstream bioinformatics and functional analyses, including clustering, data visualization, GO annotation and pathway analysis. DE genes were also subjected to Enrichr analysis (https://maayanlab.cloud/Enrichr/) for pathway analysis. Heatmaps were generated by feeding the Variance-Stabilizing Transformed values of selected DE genes from DESeq2 into Morpheus (https://software.broadinstitute.org/morpheus/).

### Quantitative real-time qRT-PCR Analysis

Total RNA was isolated using RNeasy mini kit (Qiagen #74106) with in-column genomic DNA digestion step (RNAse-free DAase set, Qiagen #79254) according to the manufacturer’s protocol. Reverse transcription was performed with 400ng of total RNA using iScript Reverse Transcription supermix (Bio-Rad #1708841) in a 20ul reaction. The cDNA mixture was diluted 1:20 and 4ul of the diluted cDNA was taken to perform qPCR using the Power SYBR green PCR master mix (Fisher Scientific #4368708) in CFX96 Touch™ Real-Time PCR Detection System (Bio-Rad). Data was normalized to internal control 18s RNA levels. Relative expression was calculated using 2-ΔΔCt method. All samples were run in technical triplicates and at least three independent experiments were performed. Primer sequences are outlined in Table 2 below.

**Table 2.**
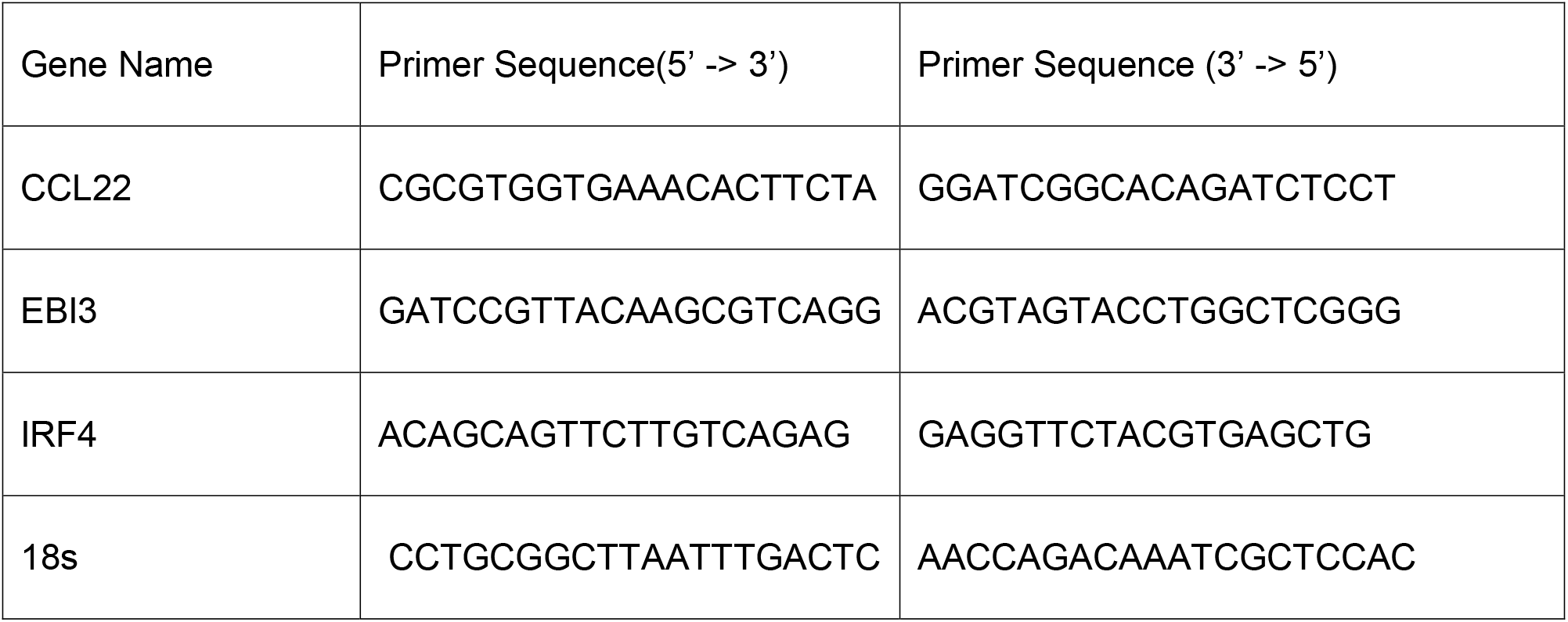
RT-PCR primers used in this study.

### Immunoblot Analysis

Cells were lysed in Laemelli buffer (0.2M Tris-HCL, 0.4 M Dithiothreitol, 277mM SDS, 6mM Bromophenol blue and 10% glycerol v/v) and sonicated at 4℃ for five seconds using a probe sonicator at 20% amplitude and boiled at 95℃ for eight minutes. The whole cell lysates were resolved by 12% or 15% SDS-PAGE, transferred to nitrocellulose filters at 100V at 4℃ for 1.5 hour, blocked with 5% non-fat dried milk in 1X TBST for 30 min at room-temperature, and then probed with the indicated primary antibodies (diluted in 1x TBS-T with 0.02% sodium azide at recommended manufacturer concentrations) overnight at 4℃ on a rotating platform. Blots were washed three times in TBST for ten minutes each, and then probed with horse-radish peroxidase (HRP)-conjugated secondary antibodies, at a dilution of 1:3000 in 1X TBST with 5% non-fat dried milk. Blots were then washed three times in TBST for ten minutes each, developed by ECL chemiluminescent substrate (Thermo Scientific, #34578) and imaged on Li-COR Odyssey workstation.

### Flow cytometry analysis

FACS was performed using a FACSCalibur instrument (BD). For ICAM-1 and FAS detection, cells were washed in PBS w/2%FBS, then stained on ice for 30 min with BioLegend PE-conjugated anti-CD54/ICAM-1 and APC-conjugated anti-CD95/Fas antibody, washed three times with PBS with 2%FBS and analyzed by FACS. 7-AAD viability assays were carried out suing 7-AAD (Thermo Fisher, #A1310) where cells were harvested and washed twice in 1XPBS supplemented with 2% FBS (Gibco). Washed cells were incubated with 1ug/ml 7-AAD solution in 1X PBS/2%FBS buffer for five minutes at room temperature and protected from light. Stained cells were analyzed via flow cytometry. Annexin V assay was performed with harvesting 1 X 10^6 cells and washed with 1X PBS twice to remove excess RPMI. 2 X10^5 cells were then resuspended to be stained with 5ul of Annexin V-FITC (#640945, Biolegend) in 100ul of Annexin V binding buffer (10mM HEPES, 140mM NaCl and 2.5mM CaCl_2_). Cells were incubated at room temperature for 15 minutes and protected from light before analyzed by FACS.

### Bioinformatic analysis and Software

All the growth curves and column charts were made with GraphPad Prism v.9. FACS data was analyzed by FlowJo V10.

## Figure Legends

**Fig. S1.**
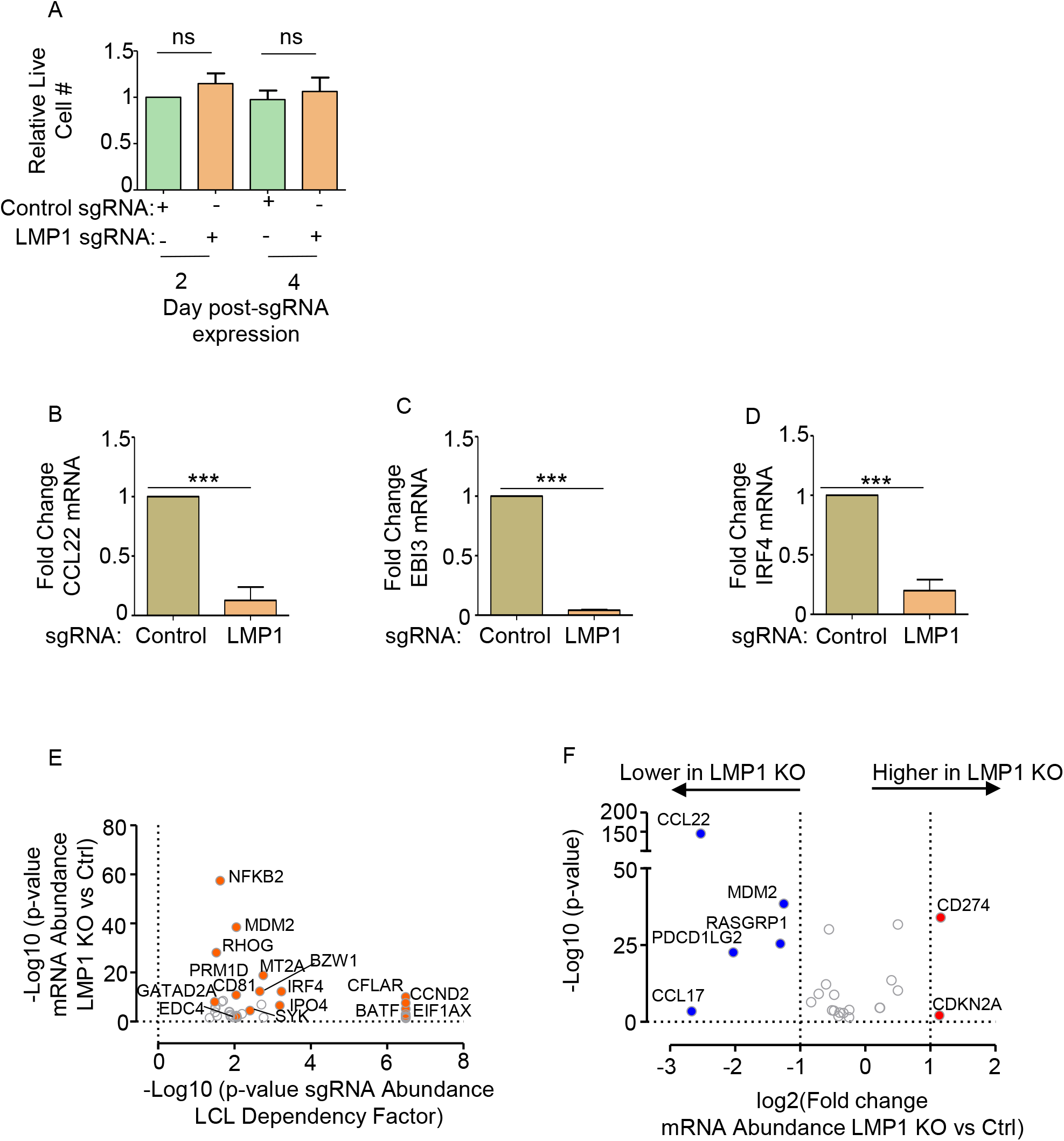
Characterization of LMP1 KO effects on GM12878 LCL target gene regulation. (A)Relative mean + standard deviation (SD) live cell numbers from CellTitreGlo analysis of n=3 replicates of Cas9+ GM12878 LCLs, transduced with lentiviruses that expressed control or LMP1 sgRNAs and puromycin selected, for 2 versus 4 days. (B) RT-PCR analysis of CCL22 mRNA abundance in Cas9+ GM12878 post-transduction with lentiviruses that expressed control or LMP1 sgRNA and puromycin selected for 2 days, as in **Fig. 1B**. The same RNA used for RNA-seq (**Fig. 1B**) was used for these qPCR experiments. Values from cells with control sgRNA were set to 1, and mean fold-change of CCL22 mRNA abundance + SD in cells with LMP1 sgRNA are shown. p-values were determined by one-sided Fisher’s exact test from two independent experiments, each with two technical replicates. ***p<0.001. (C) RT-PCR analysis of EBI3 mRNA abundance in Cas9+ GM12878 post-transduction with lentiviruses that expressed control or LMP1 sgRNA and puromycin selected for 2 days, as in **Fig. 1B**. The same RNA used for RNA-seq (**Fig. 1B**) was used for these qPCR experiments. Values from cells with control sgRNA were set to 1, and mean fold-change of EBI3 mRNA abundance + SD in cells with LMP1 sgRNA are shown. p-values were determined by one-sided Fisher’s exact test from two independent experiments, each with two technical replicates. ***p<0.001. (D) RT-PCR analysis of IRF4 mRNA abundance in Cas9+ GM12878 post-transduction with lentiviruses that expressed control or LMP1 sgRNA and puromycin selected for 2 days, as in **Fig. 1B**. The same RNA used for RNA-seq (**Fig. 1B**) was used for these qPCR experiments. Values from cells with control sgRNA were set to 1, and mean fold-change of IRF4 mRNA abundance + SD in cells with LMP1 sgRNA are shown. p-values were determined by one-sided Fisher’s exact test from two independent experiments, each with two technical replicates. ***p<0.001. (E) Scatter plot analysis cross-comparing the significance of changes in LCL dependency factor expression upon GM127878 LMP1 KO versus the CRISPR screen significance score for selection against sgRNAs in LCL vs Burkitt dependency factor analysis (25). Shown on the Y-axis are -log10 transformed P-values from RNAseq analysis of GM12878 LCLs transduced with lentiviruses expressing LMP1 versus control sgRNA (as in **Fig. 1F**), versus -log10 transformed P-values from CRISPR LCL vs Burkitt cell dependency factor analysis (25). Higher Y-axis scores indicate more significant differences in expression for the indicated genes in GM12878 with LMP1 vs control sgRNA. Higher X-axis scores indicate a stronger selection against sgRNA targeting the indicated genes in GM12878 LCLs versus P3HR1 Burkitt cells over 21 days of cell culture. Shown are genes with p<0.05 in both analyses. (F) Volcano plot analysis visualizing KEGG Hodkin lymphoma pathway gene -Log10 (P-value) on the y-axis versus log2 transformed fold change in mRNA abundances on the x-axis of GM12878 genes in cells expressing LMP1 versus control sgRNA (as in **Fig. 1F**). P-value <0.05 and >2-fold change mRNA abundance cutoffs were used.

**Fig. S2.**
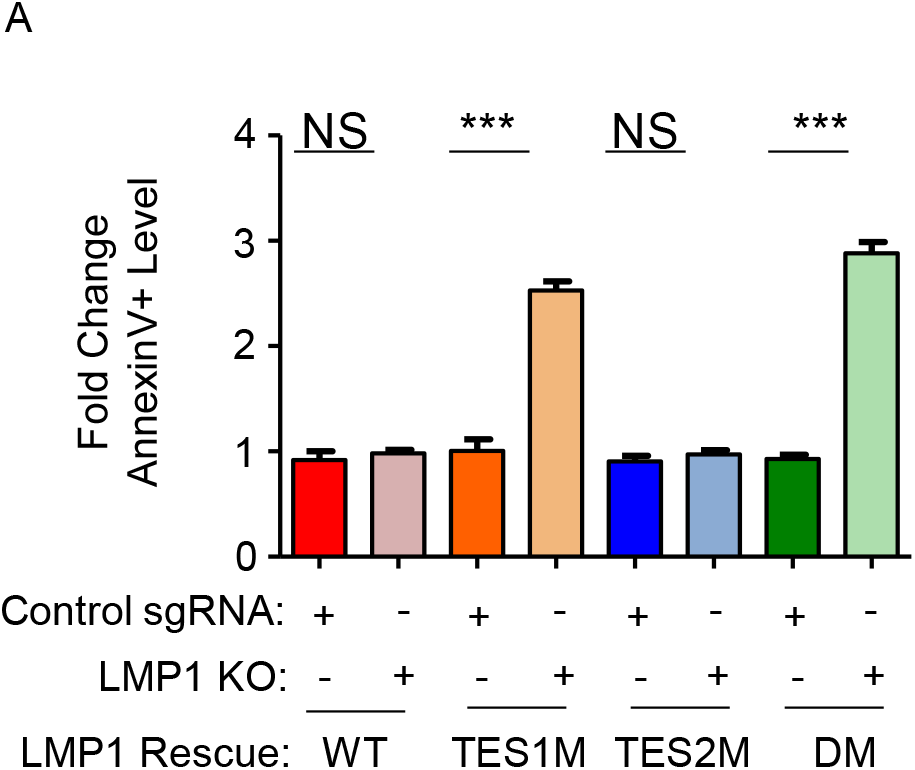
Loss of TES1 but not TES2 signaling triggers LCL apoptosis. Mean ± SD of fold change plasma membrane Annexin V values from n=3 independent experiments, using GM12878 with the indicated control or LMP1 sgRNA and rescue cDNA expression. Values in GM12878 with control sgRNA and no LMP1 rescue cDNA were set to 1.

**Figure S3.**
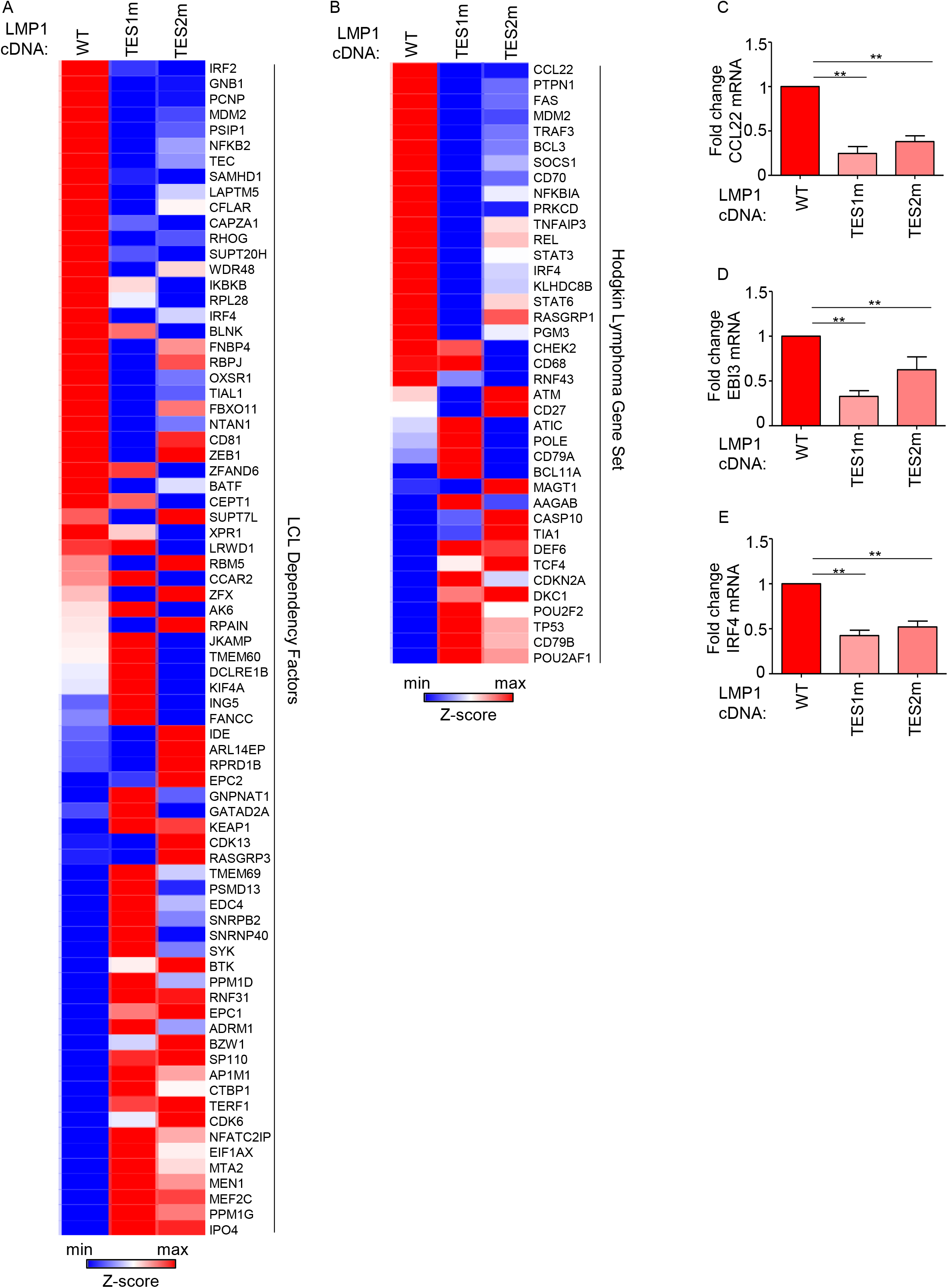
Characterization of TES1 vs TES2 LCL dependency factor and Hodgkin lymphoma pathway targets. (A) Heatmap analysis of CRISPR defined LCL dependency factor gene relative row Z-scores from RNAseq of GM12878 expressing LMP1 sgRNA and the indicated rescue cDNA, as in **Fig. 3**. The Z-score scale is shown at bottom, where blue and red colors indicate lower versus higher relative expression, respectively. Two-way ANOVA P-value cutoff of <0.05 and >2-fold gene expression cutoffs were used. (B) Heatmap analysis of KEGG Hodgkin Lymphoma pathway gene relative row Z-scores from RNAseq of GM12878 expressing LMP1 sgRNA and the indicated rescue cDNA, as in **Fig. 3**. Two-way ANOVA P-value cutoff of <0.05 and >2-fold gene expression cutoffs were used. (C) RT-PCR analysis of CCL22 mRNA abundance in GM12878 LCLs transduced with lentivirus expressing LMP1 sgRNA and induced for WT, TES1m or TES2m rescue cDNA expression for 6 days, as in **Fig. 3A**. The same RNA used for RNA-seq in **Fig. 3A** was used for these qPCR experiments. Values from cells with WT LMP1 rescue cDNA were set to 1, and mean fold-change of CCL22 mRNA abundance + SD in cells with TES1m or TES2m rescue cDNA are shown. p-values were determined by one-sided Fisher’s exact test from two independent experiments, each with two technical replicates. ***p<0.001. (D) RT-PCR analysis of EBI3 mRNA abundance in GM12878 LCLs transduced with lentivirus expressing LMP1 sgRNA and induced for WT, TES1m or TES2m rescue cDNA expression for 6 days, as in **Fig. 3A**. The same RNA used for RNA-seq in **Fig. 3A** was used for these qPCR experiments. Values from cells with WT LMP1 rescue cDNA were set to 1, and mean fold-change of EBI3 mRNA abundance + SD in cells with TES1m or TES2m rescue cDNA are shown. p-values were determined by one-sided Fisher’s exact test from two independent experiments, each with two technical replicates. ***p<0.001. (E) RT-PCR analysis of IRF4 mRNA abundance in GM12878 LCLs transduced with lentivirus expressing LMP1 sgRNA and induced for WT, TES1m or TES2m rescue cDNA expression for 6 days, as in **Fig. 3A**. The same RNA used for RNA-seq in **Fig. 3A** was used for these qPCR experiments. Values from cells with WT LMP1 rescue cDNA were set to 1, and mean fold-change of IRF4 mRNA abundance + SD in cells with TES1m or TES2m rescue cDNA are shown. p-values were determined by one-sided Fisher’s exact test from two independent experiments, each with two technical replicates. ***p<0.001.

**Fig. S4.**
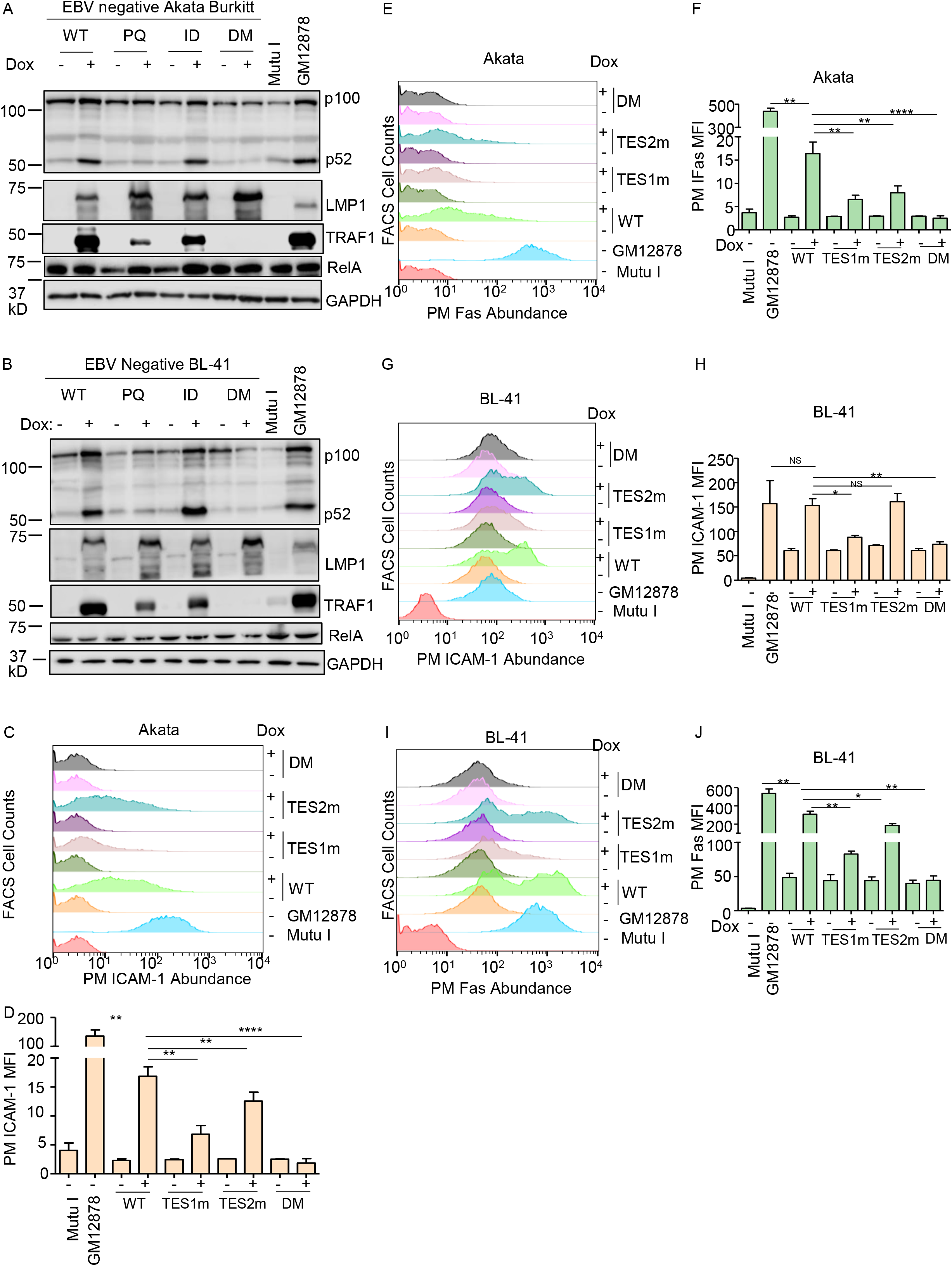
Validation of LMP1 WT, TES1m, TES2m and DM conditional expression system in EBV-negative Akata Burkitt B-cells. (A) Immunoblot analysis of WCL from Akata cells induced for LMP1 WT, TES1m, TES2m or DM expression by addition of 250 ng/ml doxycycline (Dox) for 24 hours, as indicated. For cross-comparison, WCL from equal numbers of Mutu I Burkitt lymphoma (latency I, lacks LMP1 expression) and GM12878 were also included at right. Blots are representative of n = 3 experiments. (B) Immunoblot analysis of WCL from BL-41 cells induced for WT, TES1m, TES2m or DM LMP1 by addition of 250 ng/ml Dox for 24 hours, as indicated. For cross-comparison, WCL from equal numbers of Mutu I Burkitt and GM12878 were also included at right. Blots are representative of n = 3 experiments. (C) FACS analysis of plasma membrane (PM) ICAM-1 abundance in Akata cells induced for LMP1 by 250 ng/ml Dox for 24 hours, as indicated. Y-axis are histogram cell counts, X-axis represents PM ICAM-1 abundance. For comparison, levels in GM12878 LCLs or latency I Mutu I Burkitt cells are shown. (D) PM ICAM-1 Mean Fluorescence Intensity (MFI) + standard deviation (SD) from n=3 replicates in Akata cells with the indicated LMP1 expression, as in (C). p-values were determined by one-sided Fisher’s exact test. **p<0.001, ***p<0001. (E) FACS analysis of PM Fas abundance in Akata cells induced for LMP1 by 250ng/ml of Dox for 24 hours as indicated. For comparison, GM12878 LCLs or latency I Mutu I Burkitt cells were also analyzed. (F) PM Fas MFI + SD from n=3 replicates in Akata cells with the indicated LMP1 expression, as in (E). P-values were determined by one-sided Fisher’s exact test. **p<0.001, ***p<0.0001. (G) FACS analysis of PM ICAM-1 in BL-41 cells induced for LMP1 expression by 250ng/ml Dox for 24 hours, as indicated. For comparison, GM12878 and Mutu I were also analyzed. p-values were determined by one-sided Fisher’s exact test. **p<0.001, ***p<0.0001. (H) PM ICAM-1 MFI + SD from n=3 replicates in BL-41 cells with the indicated LMP1 expression, as in (G). P-values were determined by one-sided Fisher’s exact test. **p<0.001, ***p<0.0001. (I) FACS analysis of PM Fas levels in BL-41 cells induced for LMP1 expression by 250 ng/ml dox for 24 hours, as indicated. GM12878 and Mutu I were analyzed for cross-comparison. p-values were determined by one-sided Fisher’s exact test. **p<0.001, ***p<0.0001. (J) PM Fas MFI + SD from n=3 replicates of BL-41-LMP1 with the indicated LMP1 expression, as in (I). Mutu I and GM12878 were analyzed for comparison. p-values were determined by one-sided Fisher’s exact test. **p<0.001, ***p<0.0001.

**Fig. S5.**
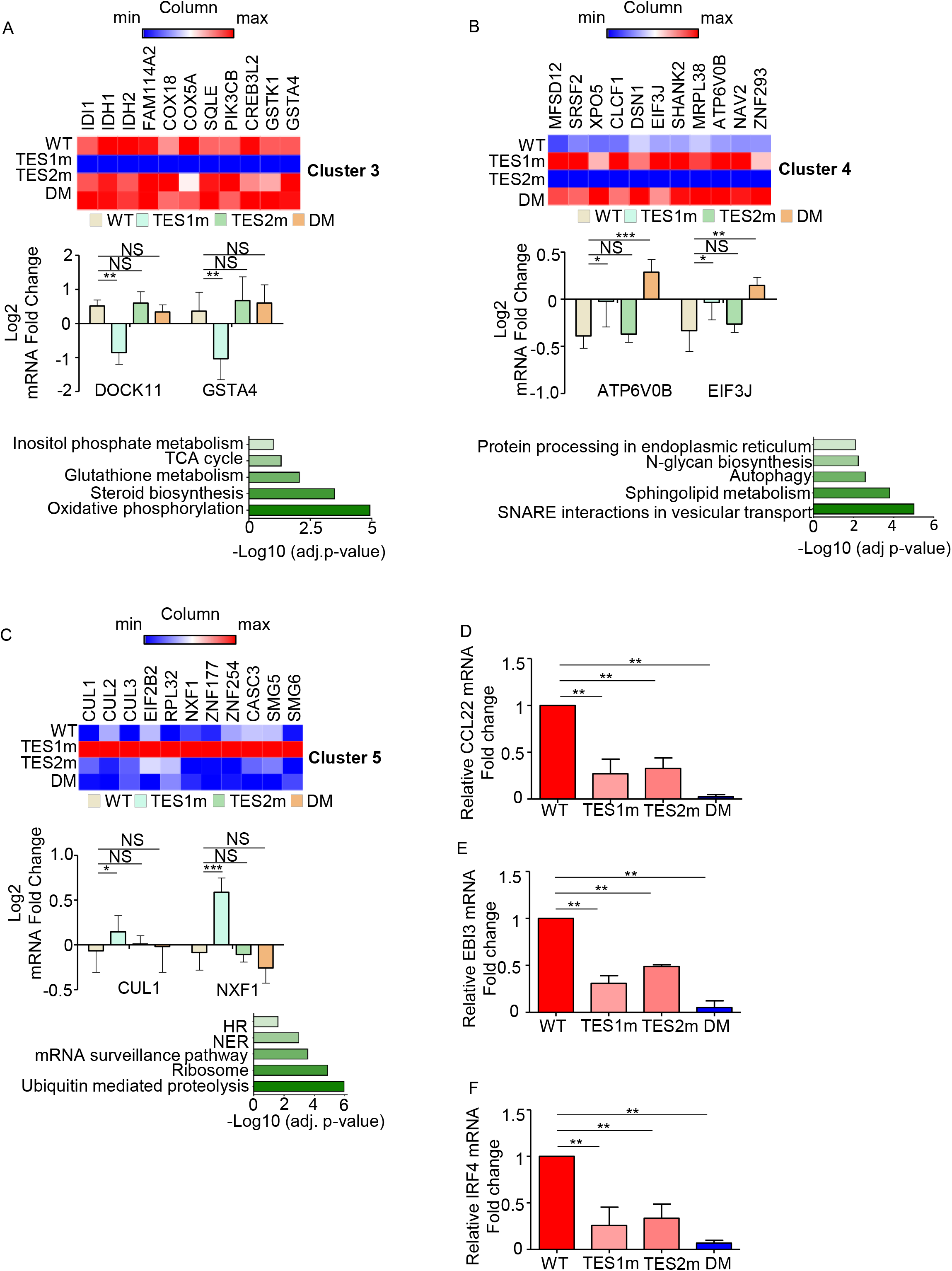
Characterization of host genome-wide Akata B-cell LMP1 target genes, related to Figure 4. (A) Heatmaps of representative **Figure 4** Cluster 3 differentially regulated genes (top), with column maximum colored red and minimum colored blue, as shown by the scalebar. Also shown are expression values of two representative Clusters 3 genes (lower left) and Enrichr analysis of KEGG pathways most significantly enriched Cluster 3 gene sets (lower right). p-values were determined by one-sided Fisher’s exact test. **p<0.01. (B) Heatmaps of representative **Figure 4** Cluster 4 differentially regulated genes (top), with column maximum colored red and minimum colored blue, as shown by the scalebar. Also shown are expression values of two representative Clusters 4 genes (lower left) and Enrichr analysis of KEGG pathways most significantly enriched Cluster 4 gene sets (lower right). p-values were determined by one-sided Fisher’s exact test. *p<0.05, **p<0.01, ***p<0.001. (C) Heatmaps of representative **Figure 4** Cluster 5 differentially regulated genes (top), with column maximum colored red and minimum colored blue, as shown by the scalebar. Also shown are expression values of two representative Clusters 5 genes (lower left) and Enrichr analysis of KEGG pathways most significantly enriched Cluster 5 gene sets (lower right). p-values were determined by one-sided Fisher’s exact test. *p<0.05, ***p<0.001. (D) RT-PCR analysis of CCL22 mRNA abundance in cells with conditional LMP1 WT, TES1m, TES2m or DM expression induced by 250 ng/ml doxycycline for 24 hours as in Fig. 5A. The same RNA used for RNA-seq in Fig. 5A was used for these qPCR experiments. Values from cells with conditional WT LMP1 expression were set to 1, and mean fold-change of CCL22 mRNA abundance + SD in cells with conditional TES1m,TES2m or DM expression are shown. p-values were determined by one-sided Fisher’s exact test from two independent experiments, each with two technical replicates. **p<0.01. (E) RT-PCR analysis of EBI3 mRNA abundance in cells with conditional LMP1 WT, TES1m, TES2m or DM expression induced by 250 ng/ml doxycycline for 24 hours as in Fig. 5A. The same RNA used for RNA-seq in Fig. 5A was used for these qPCR experiments. Values from cells with conditional WT LMP1 expression were set to 1, and mean fold-change of CCL22 mRNA abundance + SD in cells with conditional TES1m,TES2m or DM expression are shown. p-values were determined by one-sided Fisher’s exact test from two independent experiments, each with two technical replicates. **p<0.01. (F) RT-PCR analysis of IRF4 mRNA abundance in cells with conditional LMP1 WT, TES1m, TES2m or DM expression induced by 250 ng/ml doxycycline for 24 hours as in Fig. 5A. The same RNA used for RNA-seq in Fig. 5A was used for these qPCR experiments. Values from cells with conditional WT LMP1 expression were set to 1, and mean fold-change of CCL22 mRNA abundance + SD in cells with conditional TES1m,TES2m or DM expression are shown. p-values were determined by one-sided Fisher’s exact test from two independent experiments, each with two technical replicates. **p<0.01.

**Fig. S6.**
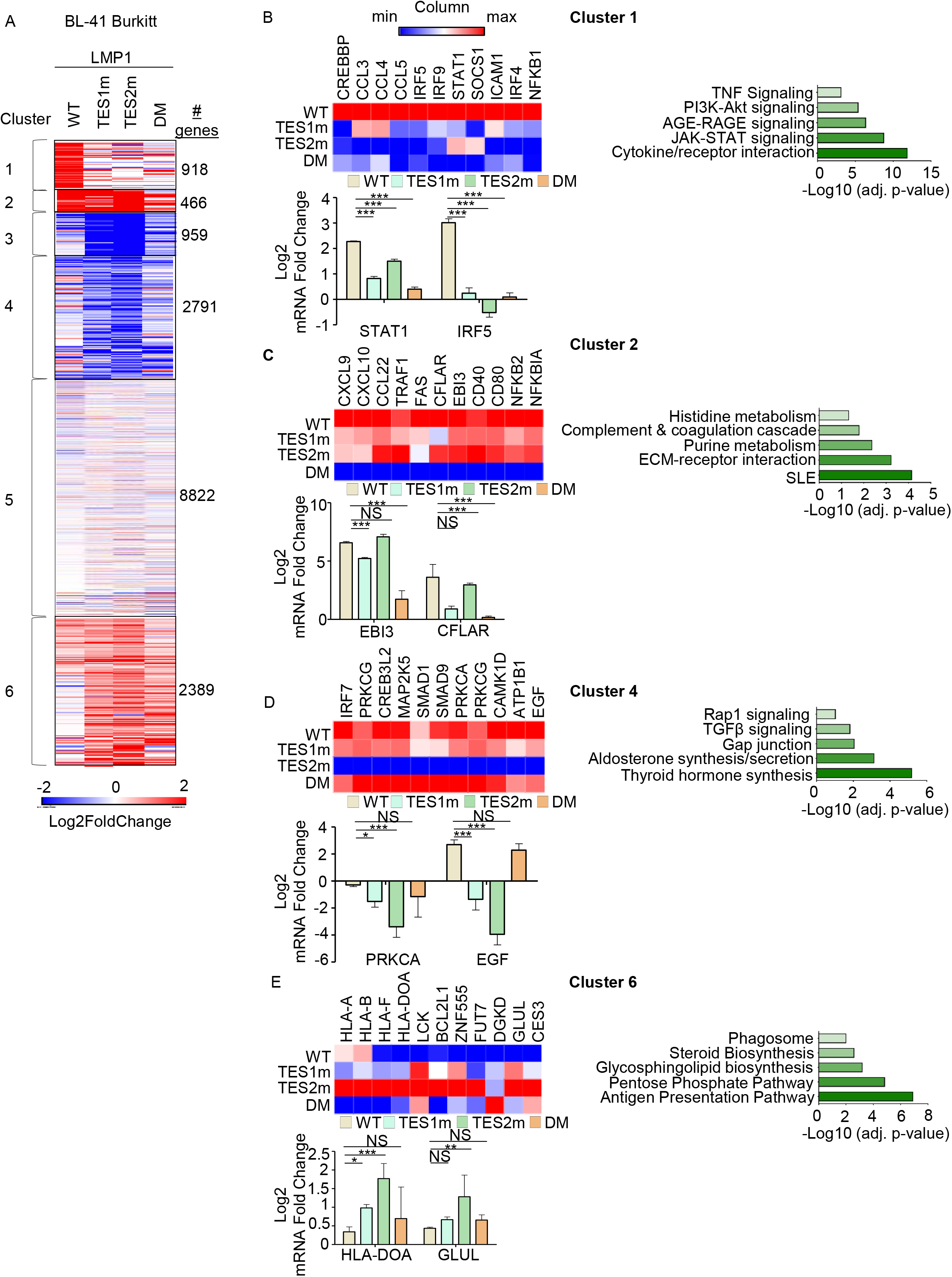
RNAseq analysis of BL-41 B-cell responses to WT, TES1, TES2 or DM LMP1. (A) K-means heatmap analysis of RNAseq datasets from n=3 replicates generated in EBV-BL-41 Burkitt cells with conditional LMP1 WT, TES1m, TES2m or DM expression induced by 250 ng/ml doxycycline for 24 hours. The heatmap visualizes host gene Log2 Fold change across the four conditions, divided into six clusters. A two-way ANOVA P value cutoff of <0.01 and >2-fold gene expression were used. # of genes in each cluster is indicated at right. (B) Heatmaps of representative Cluster 1 differentially regulated genes (top), with column maximum (max) colored red and minimum (min) colored blue, as shown by the scalebar. Also shown are expression values of two representative Cluster 1 genes (lower left) and Enrichr analysis of KEGG pathways significantly enriched in Cluster 1 gene sets (lower right). p-values were determined by one-sided Fisher’s exact test. ***p<0.001. (C) Heatmaps of representative Cluster 2 differentially regulated genes (top). Also shown are expression values of two representative Cluster 2 genes (lower left) and Enrichr analysis of KEGG pathways significantly enriched in Cluster 2 gene sets (lower right). p-values were determined by one-sided Fisher’s exact test. ***p<0.001. (D) Heatmaps of representative Cluster 4 differentially regulated genes (top). Also shown are expression values of two representative Cluster 4 genes (lower left) and Enrichr analysis of KEGG pathways significantly enriched in Cluster 4 gene sets (lower right). p-values were determined by one-sided Fisher’s exact test. *p<0.05, ***p<0.001. (E) Heatmaps of representative Cluster 6 differentially regulated genes (top). Also shown are expression values of two representative Cluster 6 genes (lower left) and Enrichr analysis of KEGG pathways significantly enriched in Cluster 6 gene sets (lower right). p-values were determined by one-sided Fisher’s exact test.*p<0.05, **p<0.01, ***p<0.001.

**Fig. S7.**
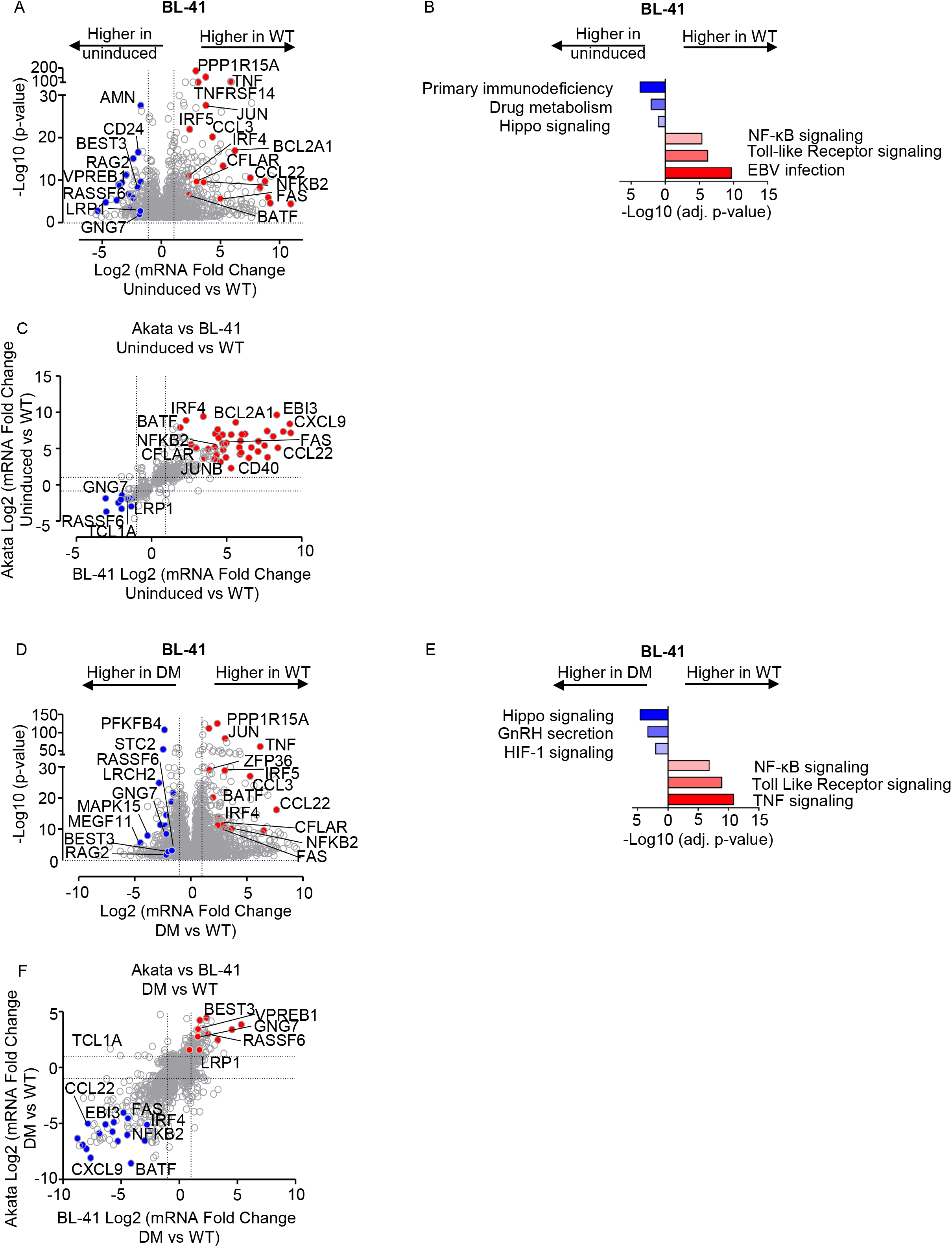
Cross-comparison of WT and DM LMP1 effects on Akata vs BL-41 transcriptomes. (A) Volcano plot analysis of host transcriptome-wide genes differentially expressed in BL-41 cells conditionally induced for WT LMP1 expression for 24h by 250 ng/ml Dox versus in mock induced cells. Higher X-axis fold changes indicate genes more highly expressed in cells with WT LMP1 expression, whereas lower X-axis fold changes indicate higher expression in cells mock induced for LMP1. Data are from n=3 RNAseq datasets. (B) Enrichr analysis of KEGG pathways most highly enriched in RNAseq data as in (A) amongst genes more highly expressed in BL-41 with WT LMP1 (red) vs amongst genes more highly expressed with mock LMP1 induction (blue). (C) Volcano plot cross-comparison of Log2 transformed fold change of host mRNA levels in BL-41 cells (X-axis) versus Akata cells (Y-axis) uninduced versus induced for WT LMP1 by 250 ng/ml Dox for 24 hours. Selected genes highly WT LMP1 induced in both Burkitt contexts are highlighted in red, whereas selected genes suppressed by LMP1 in both Burkitt contexts are highlighted in blue. (D) Volcano plot analysis of host transcriptome-wide genes differentially expressed in BL-41 cells conditionally induced for DM versus WT LMP1 expression for 24h by 250 ng/ml Dox. Higher X-axis fold changes indicate genes more highly expressed in cells with WT LMP1 expression, whereas lower X-axis fold changes indicate higher expression in cells induced for DM LMP1. Data are from n=3 RNAseq datasets. (E) Enrichr analysis of KEGG pathways most highly enriched in RNAseq data as in (D) amongst genes more highly expressed in BL-41 with WT LMP1 (red) vs amongst genes more highly expressed with DM LMP1 induction (blue). (F) Volcano plot cross-comparison of Log2 transformed fold change of host mRNA levels in BL-41 cells (X-axis) versus Akata cells (Y-axis) induced for DM versus WT LMP1 by 250 ng/ml Dox for 24 hours. Selected genes highly WT LMP1 induced in both Burkitt contexts relative to levels in cells with DM LMP1 expression are highlighted in red, whereas selected genes suppressed by WT LMP1 in both Burkitt contexts are highlighted in blue.

**Fig. S8.**
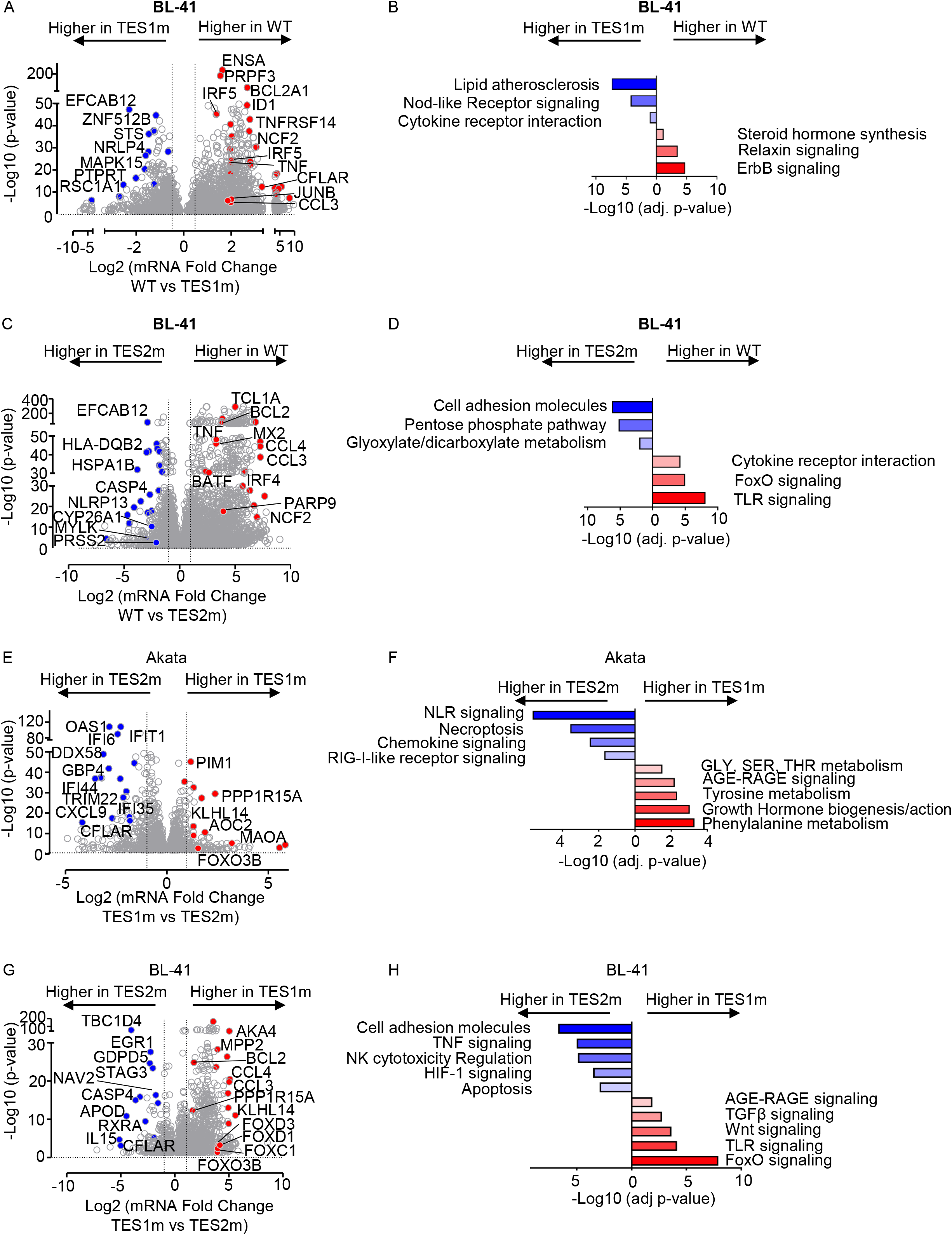
Cross-comparison of TES1 and TES2 LMP1 effects on Akata vs BL-41 transcriptomes. A) Volcano plot analysis of host transcriptome-wide genes differentially expressed in BL-41 cells conditionally induced for TES1m vs WT LMP1 expression for 24h by 250 ng/ml Dox. Higher X-axis fold changes indicate genes more highly expressed in cells with WT LMP1 expression, whereas lower X-axis fold changes indicate higher expression induced for TES1m LMP1. Data are from n=3 RNAseq datasets. (B) Enrichr analysis of KEGG pathways most highly enriched in RNAseq data as in (A) amongst genes more highly expressed in BL-41 with WT LMP1 (red) vs amongst genes more highly expressed with TES1m LMP1 induction (blue). (C) Volcano plot analysis of host transcriptome-wide genes differentially expressed in BL-41 cells conditionally induced for TES2m vs WT LMP1 expression for 24h by 250 ng/ml Dox. Higher X-axis fold changes indicate genes more highly expressed in cells with WT LMP1 expression, whereas lower X-axis fold changes indicate higher expression induced for TES2m LMP1. Data are from n=3 RNAseq datasets. (D) Enrichr analysis of KEGG pathways most highly enriched in RNAseq data as in (A) amongst genes more highly expressed in BL-41 with WT LMP1 (red) vs amongst genes more highly expressed with TES2m LMP1 induction (blue). (E) Volcano plot analysis of host transcriptome-wide genes differentially expressed in Akata cells conditionally induced for TES1m vs TES2m LMP1 expression for 24h by 250 ng/ml Dox. Higher X-axis fold changes indicate genes more highly expressed in cells with TES1m LMP1 expression, whereas lower X-axis fold changes indicate higher expression induced for TES2m LMP1. Data are from n=3 RNAseq datasets. (F) Enrichr analysis of KEGG pathways most highly enriched in RNAseq data as in (E) amongst genes more highly expressed in Akata with TES1m LMP1 (red) vs amongst genes more highly expressed with TES2m LMP1 induction (blue). (G) Volcano plot analysis of host transcriptome-wide genes differentially expressed in BL-41 cells conditionally induced for TES1m vs TES2m LMP1 expression for 24h by 250 ng/ml Dox. Higher X-axis fold changes indicate genes more highly expressed in cells with TES1m LMP1 expression, whereas lower X-axis fold changes indicate higher expression induced for TES2m LMP1. Data are from n=3 RNAseq datasets. (H) Enrichr analysis of KEGG pathways most highly enriched in RNAseq data as in (G) amongst genes more highly expressed in BL-41 with TES1m LMP1 (red) vs amongst genes more highly expressed with TES2m LMP1 induction (blue).

**Figure S9.**
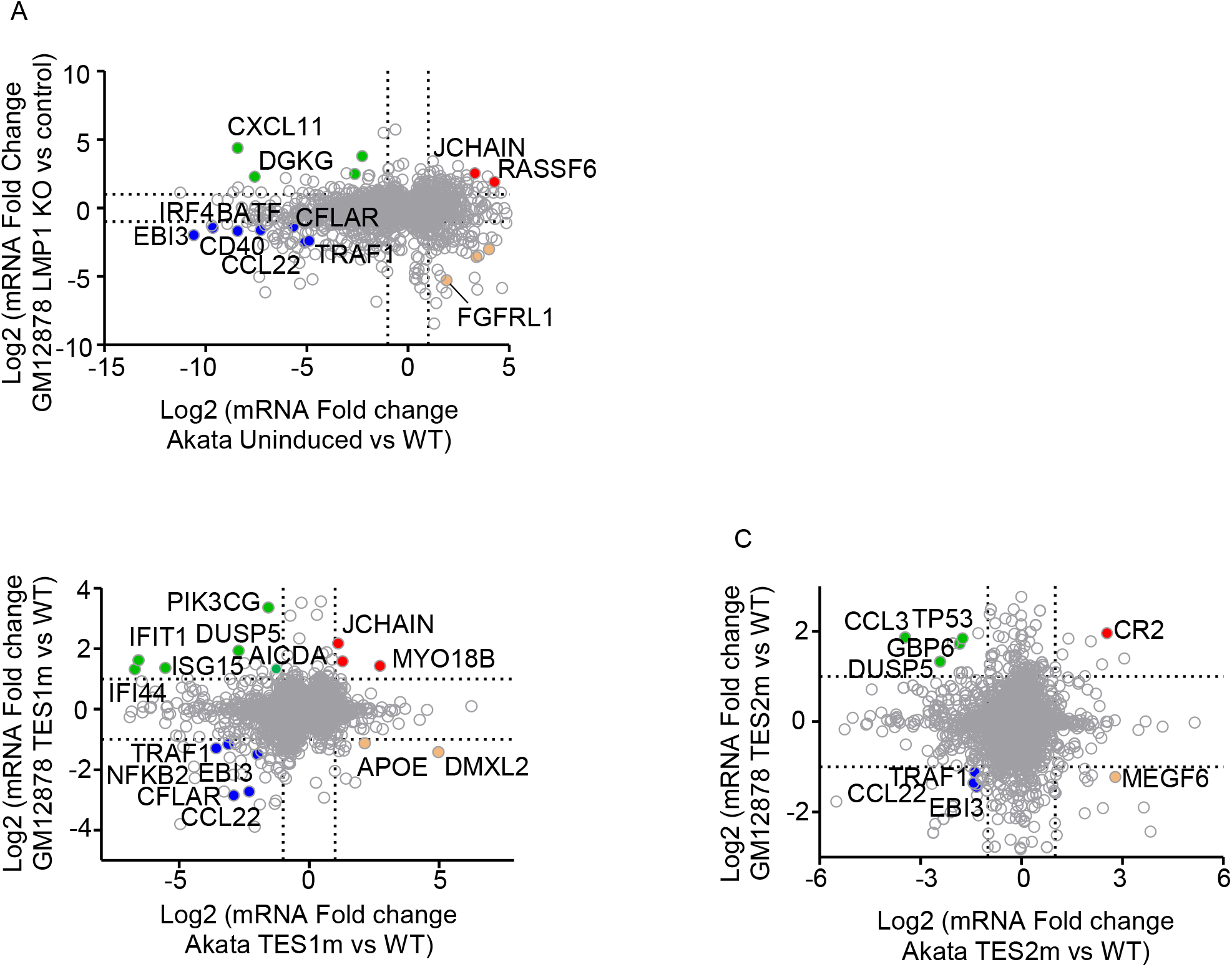
Cross-comparison of host genes differentially expressed upon perturbation of LCL LMP1 versus upon LMP1 induction in Akata cells. (A) Volcano plot analysis of host genes differentially expressed upon WT LMP1 induction in Akata (X-axis) versus upon LMP1 KO in GM12878 (Y-axis). Shown are Log2 transformed mRNA fold change values for Akata cells mock induced versus induced for LMP1 WT expression for 24 hours (X-axis) versus upon expression of LMP1 vs control sgRNA in GM18278 for 48 hours. Genes more highly expressed in mock-induced Akata have higher x-axis values, whereas genes more highly expressed in Akata induced for WT LMP1 have lower x-axis values. Likewise, genes with higher expression with control sgRNA expression have higher y-axis values, whereas genes with lower expression in GM12878 with LMP1 KO have lower Y-axis values. P value <0.05 and >2-fold gene expression cutoffs were used. (B) Volcano plot analysis of host genes differentially expressed upon TES1m vs WT LMP1 induction in Akata (X-axis) versus upon rescue of LMP1 KO GM12878 with TES1m versus WT LMP1 (Y-axis). Shown are Log2 transformed mRNA fold change values for Akata cells induced for TES1m versus WT LMP1 expression for 24 hours (X-axis) versus upon rescue of GM18278 LMP1 KO with TES1m vs WT LMP1 cDNA, as in **Fig 3**. Genes more highly expressed in Akata with TES1m than WT LMP1 expression have higher x-axis values, whereas genes more highly expressed in Akata induced for WT LMP1 than TES1m have lower x-axis values. Likewise, GM12878 genes with higher expression with TES1m rescue have higher y-axis values, whereas genes with lower expression in GM12878 with TES1m than WT rescue have lower Y-axis values. P value <0.05 and >2-fold gene expression cutoffs were used. (C) Volcano plot analysis of host genes differentially expressed upon TES2m vs WT LMP1 induction in Akata (X-axis) versus upon rescue of LMP1 KO GM12878 with TES2m versus WT LMP1 (Y-axis). Shown are Log2 transformed mRNA fold change values for Akata cells induced for TES2m versus WT LMP1 expression for 24 hours (X-axis) versus upon rescue of GM18278 LMP1 KO with TES2m vs WT LMP1 cDNA, as in **Fig 3**. Genes more highly expressed in Akata with TES2m than WT LMP1 expression have higher x-axis values, whereas genes more highly expressed in Akata induced for WT LMP1 than TES2m have lower x-axis values. Likewise, GM12878 genes with higher expression with TES2m rescue have higher y-axis values, whereas genes with lower expression in GM12878 with TES2m than WT rescue have lower Y-axis values. P value <0.05 and >2-fold gene expression cutoffs were used.

**Fig. S10.**
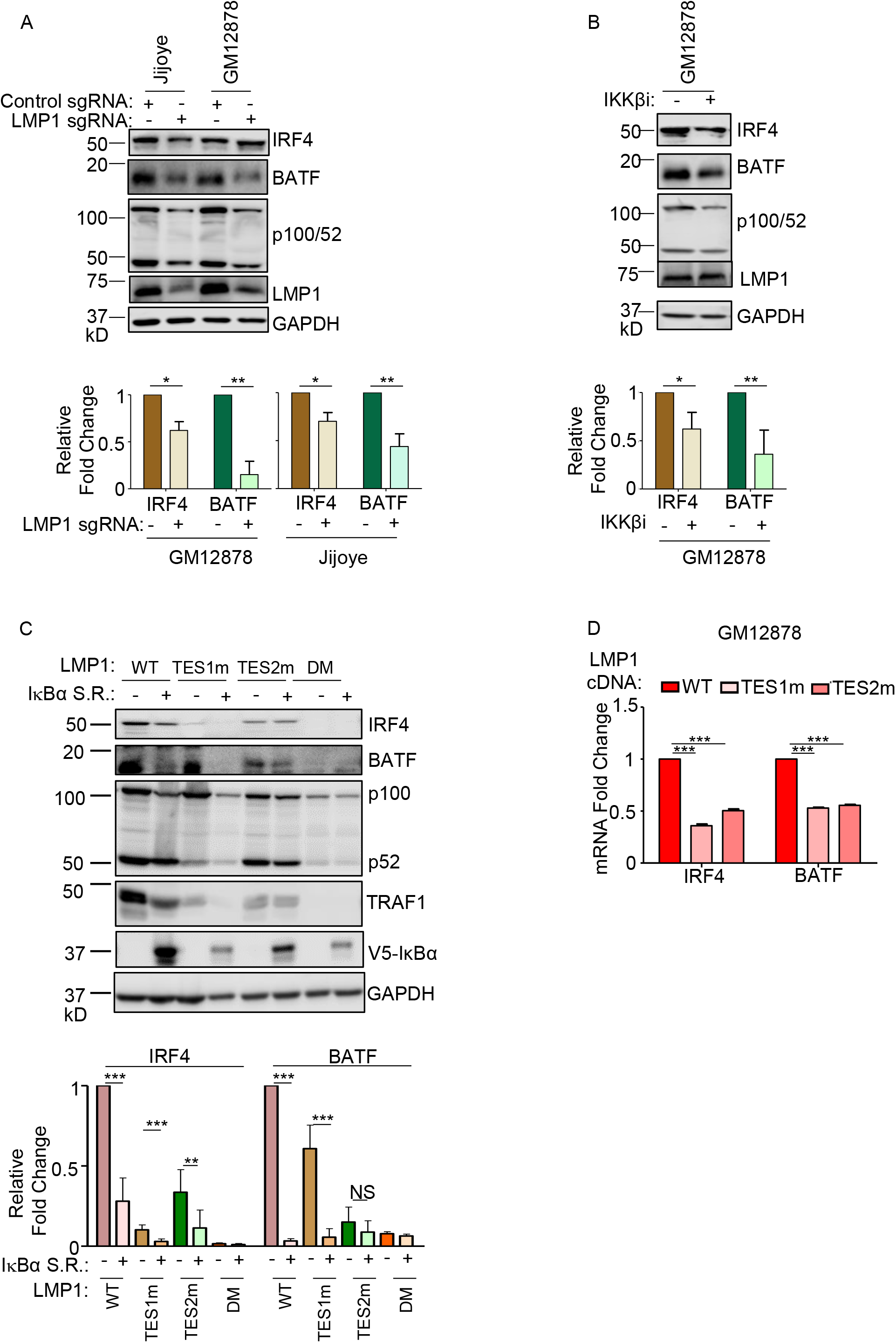
Roles of TES1 and TES2 canonical NF-κB pathways in BATF and IRF4 induction. (A) Immunoblot analysis of WCL from latency III Jijoye Burkitt cells or GM12878 LCL expressing LMP1 targeting sgRNA, as indicated. Shown below are relative fold changes + SD from n=3 replicates of IRF4 or BATF vs GAPDH load control densitometry values, with values in sgRNA control expressing cells set to 1. P-values were determined by one-sided Fisher’s exact test. **p<0.001, ***p<0.0001. (B) Relative +SD BATF and IRF4 mRNA levels from n=3 RNAseq replicates from LMP1 KO GM12878 with TES1m, TES2m or WT LMP1 rescue cDNA expression, as in **Fig 3-4**. BATF or IRF4 levels in cells with WT LMP1 rescue were defined as 1. ***p<0.0001. (C) Immunoblot analysis of WCL from GM12878 LCLs treated with vehicle control or 1 μM IKKβ inhibitor VIII for 24 hours. Shown below are relative foldchanges + SD from n=3 replicates of IRF4 or BATF vs GAPDH load control densitometry values. Levels in vehicle control treated WT LMP1 expressing cells were set to 1. P-values were determined by one-sided Fisher’s exact test. **p<0.001, ***p<0.0001. (C) Immunoblot analysis of WCL from Akata cells induced for the indicated LMP1 construct expression, either without or together with an IκBα super-repressor (IκBα-S.R.) that blocks canonical NF-κB signaling. Shown below are relative foldchanges + SD of IRF4 or BATF vs GAPDH load control densitometry values from n=3 replicates, with values in cells expressing WT LMP1 but not IκBα-set to 1. P-values were determined by one-sided Fisher’s exact test. **p<0.001, ***p<0.0001.

**Fig. S11.**
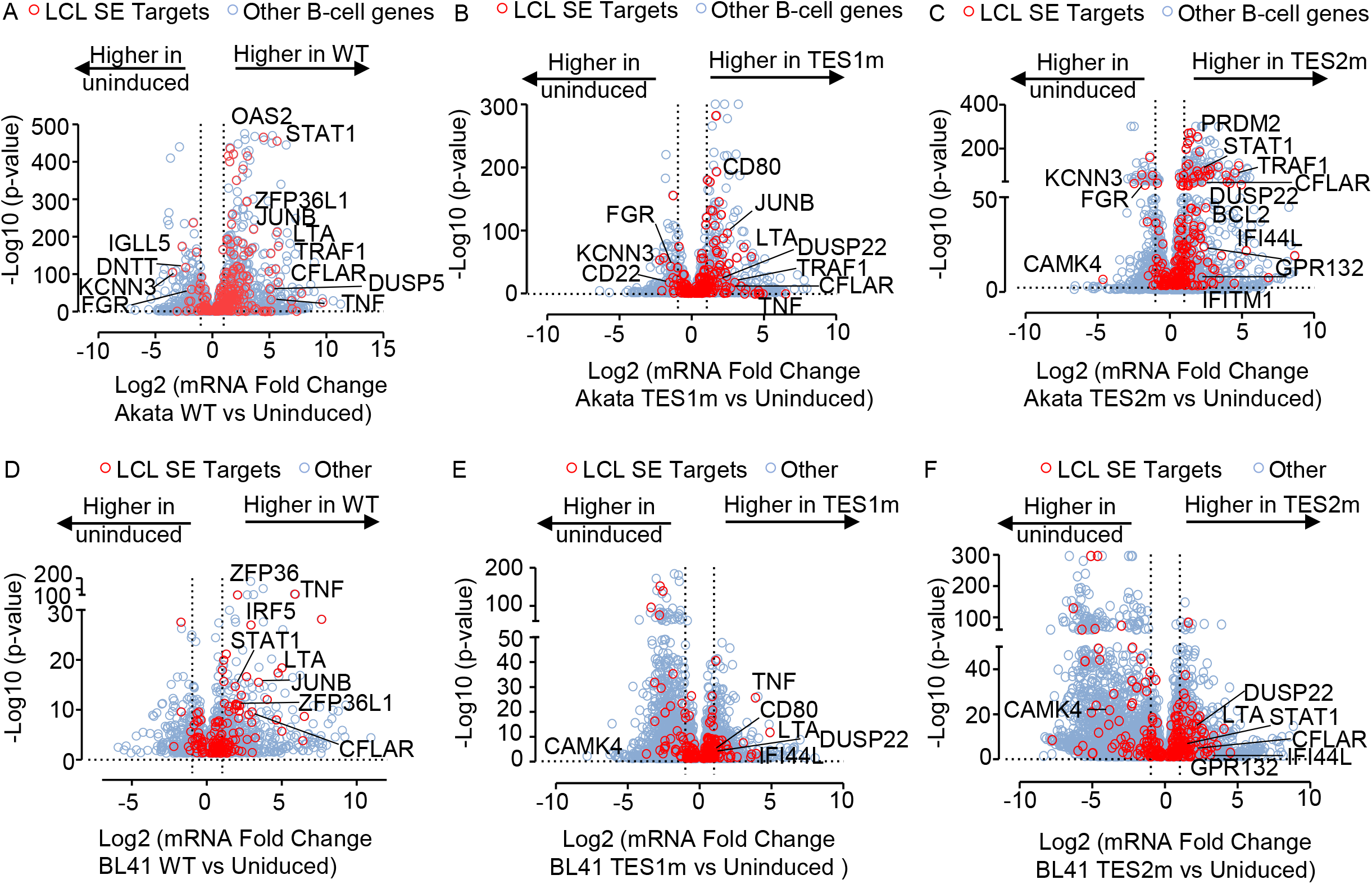
LMP1 TES1 and TES2 roles in EBV super-enhancer target gene regulation in Akata and BL-41 Burkitt B-cells. (A) Volcano plot analysis of Akata RNA-seq, comparing mRNA values in cells induced for LMP1 WT vs. mock-induced for 24 hours. SE targeted genes highlighted by red circles and other B-cell genes indicated by blue circles. P value <0.05 and >2-fold gene expression cutoffs were used. (B) Volcano plot analysis of Akata RNA-seq, comparing mRNA values in cells induced for LMP1 TES1m vs. mock-induced for 24 hours. SE targeted genes highlighted by red circles and other B-cell genes indicated by blue circles. P value <0.05 and >2-fold gene expression cutoffs were used. (C) Volcano plot analysis of Akata RNA-seq, comparing mRNA values in cells induced for LMP1 TES2m vs. mock-induced for 24 hours. SE targeted genes highlighted by red circles and other B-cell genes indicated by blue circles. P value <0.05 and >2-fold gene expression cutoffs were used. (D) Volcano plot analysis of BL-41 RNA-seq, comparing mRNA values in cells induced for LMP1 WT vs. mock-induced for 24 hours. SE targeted genes highlighted by red circles and other B-cell genes indicated by blue circles. P value <0.05 and >2-fold gene expression cutoffs were used. (E) Volcano plot analysis of BL-41 RNA-seq, comparing mRNA values in cells induced for LMP1 TES1m vs. mock-induced for 24 hours. SE targeted genes highlighted by red circles and other B-cell genes indicated by blue circles. P value <0.05 and >2-fold gene expression cutoffs were used. (F) Volcano plot analysis of BL-41 RNA-seq, comparing mRNA values in cells induced for LMP1 TES2m vs. mock-induced for 24 hours. SE targeted genes highlighted by red circles and other B-cell genes indicated by blue circles. P value <0.05 and >2-fold gene expression cutoffs were used.

**Fig. S12.** Model highlighting different modes of LMP1 TES1 and TES2 cross-talk in Burkitt B-cell target gene regulation. We identify 6 different modes of TES 1/2 cross-talk where (a) TES1 is required for the transcription of these subset of genes over TES2 (in limited amounts) or (b) TES1 or TES2 are individually sufficient for the expression of these subset of genes, or (c) TES1 is required to block the TES2-mediated suppression of this subset, or (d) TES2 is required to block the TES1-mediated suppression of this subset, or (e) TES1 is required to block the TES2-mediated upregulation of this subset, or (f) TES1 and TES2 are required for LMP1-mediated suppression of this gene subset.

